# Circular Data Analysis for Spatial Omics

**DOI:** 10.64898/2026.09.29.755467

**Authors:** Jungmin Shin, Jinkyung Yoo, Yanghyeon Cho, Arkobrato Gupta, Pin Hsun Mao, Kairavee Thakkar, Tharun Amir Eddy, Kyeong Joo Jung, Joanne Kim, Hyeongseon Jeon, Juan Xie, Dongjun Chung

## Abstract

Many biological quantities in omics are inherently periodic or directional, including circadian phase, cell-cycle position, and cellular orientation. Treating such quantities as ordinary linear variables can introduce artificial discontinuities and obscure biological interpretation. Circular methods are well developed across directional statistics, bioinformatics, and spatial statistics, but practical guidance for omics applications remains limited. We present key considerations for circular data analysis in single-cell and spatial omics, including the provenance and observational unit of the circular quantity, the inferential or predictive objective, the distinction between tissue composition and residual spatial dependence, and the choice of appropriate models. We describe relevant approaches for circular representation, inference, clustering, regression, machine learning, and spatial analysis, together with representative R software. Their practical use is illustrated through spatial transcriptomic case studies of derived circadian peak phase in a mouse model of Alzheimer’s disease and latent cell-cycle position in human ovarian cancer. These examples demonstrate how the definition of the angle, its observational unit, and tissue heterogeneity shape appropriate analysis and interpretation, including when spatial modeling is not warranted. We conclude with practical recommendations and directions for future methodological development.

## 1 Introduction

High-throughput omics technologies generate high-dimensional measurements across molecular, cellular, and spatial scales. While most molecular measurements are analyzed on ordinary linear or Euclidean scales, many biological processes are inherently periodic or directional. Circadian rhythms follow a 24-hour cycle, cell-cycle progression traverses recurrent cellular states, and tissue organization often involves orientation relative to anatomical axes or local structural features (Anafi et al.; 2017; Zheng et al.; 2022; Giese et al.; 2025). In these settings, biologically meaningful quantities are naturally represented on a circle rather than on a linear scale. Treating such quantities as ordinary numerical variables can introduce artificial discontinuities and obscure biological interpretation (Lee; 2010; Cremers and Klugkist; 2018), making circular methods increasingly essential for modern omics analyses.

Circular quantities in omics arise through multiple pathways, which we classify into direct, derived, and latent circular quantities. Some are directly observed, such as cellular orientation or physical directional measurements (Giese et al.; 2025). Others are derived from measured features, for example through polarity vectors or temporal annotations (Nelson et al.; 1979; Yang and Su; 2010; Giese et al.; 2025). Still others are latent variables inferred from high-dimensional molecular profiles, including cell-cycle position and circadian phase (Hughey et al.; 2016; Anafi et al.; 2017; Liang et al.; 2020; Zheng et al.; 2022; Auerbach et al.; 2022). Although these quantities share a common circular geometry, they differ substantially in their provenance, uncertainty, and biological interpretation. Recognizing these distinctions is essential for selecting appropriate downstream analyses and interpreting the resulting biological conclusions.

A substantial methodological foundation for circular data already exists across several disciplines. Directional statistics provides probability models, hypothesis tests, regression methods, and clustering procedures for circular observations (Fisher; 1993; Mardia and Jupp; 2000; Jammalamadaka and SenGupta; 2001; Pewsey and García-Portugués; 2021). Bioinformatics offers algorithms that infer latent circular quantities from molecular measurements (Anafi et al.; 2017; Zheng et al.; 2022; Auerbach et al.; 2022), while spatial statistics provides models for circular outcomes exhibiting spatial dependence (Jona-Lasinio et al.; 2012; Wang and Gelfand; 2014; Jona Lasinio et al.; 2020). These methods, however, are often presented as separate analytical tasks. Practical guidance is therefore limited on how to define an omics-derived circular quantity, choose an analysis according to the scientific objective, and connect inference, prediction, and spatial analysis in a biologically meaningful way.

These challenges are particularly evident in spatial omics, where complex tissue architectures introduce unique statistical hurdles. The observational unit is often distinct from the biological replication unit, complicating valid statistical inference (Rivest and Kato; 2019; Landler et al.; 2020). More importantly, tissues are intrinsically heterogeneous. Differences in cell-type composition, anatomical structure, or pathological context can generate apparent circular patterns that may be mistaken for genuine spatial dependence (Jona-Lasinio et al.; 2012; Wang and Gelfand; 2014). Distinguishing compositional effects from residual spatial structure is therefore a central prerequisite for spatial circular analysis. To address this, we advocate a composition-before-covariance strategy, ensuring that local compositional heterogeneity is rigorously accounted for before modeling residual spatial correlations.

In this paper, we present practical considerations for circular data analysis in single-cell and spatial omics. We organize them around the provenance and observational unit of the circular quantity, the inferential or predictive objective, the distinction between tissue composition and residual spatial dependence, and the choice of predictive or spatial circular models. We describe relevant representations, statistical models, and R software and illustrate their use through two spatial transcriptomic case studies involving derived circadian peak phase in a mouse model of Alzheimer’s disease (AD, Gelber et al.; 2026) and latent cell-cycle position in human ovarian cancer (10x Genomics; 2024; Zheng et al.; 2022).

The remainder of the paper follows these considerations. Section 2 introduces foundations of circular representation and inference. Section 3 reviews machine-learning approaches involving circular variables. Section 4 develops the composition-before-covariance principle for spatial modeling. Finally, Sections 5 and 6 present the examples and summarize practical considerations for future research. Reproducible R code for all analyses is provided in the Supplementary Note.

## 2 Defining and characterizing circular quantities

For analysis unit *i*, let *{***y***_i_,* **s***_i_, θ_i_,* **z***_i_, r_i_}* denote molecular measurements, spatial location, a circular quantity *θ_i_ ∈* [0, 2*π*), additional covariates, and the biological replicate from which the unit is obtained, respectively. The analysis unit may be a cell, spot, sample, or gene-by-context summary. We distinguish the analysis unit from the biological replication unit because the number of angular observations can be substantially larger than the number of independent specimens.

### 2.1 Provenance, observational unit, and representation

Circular analysis begins by defining the quantity that carries the angle, including what it represents and how it is obtained, the level at which it is observed and replicated, the reference frame in which it is defined, and how it is represented. By provenance, we distinguish directly observed, derived, and latent circular quantities. Derived and latent angles also inherit estimation uncertainty and filtering. The observational unit carrying *θ_i_* need not coincide with the biological replicate *r_i_*, and spatial analysis requires *θ_i_*and **s***_i_* to be defined on the same unit (Section 4).

An angle is interpretable only relative to a reference frame, defined by its period, angular origin, and orientation. For example, derived circadian peak phase may be anchored to Zeitgeber time (ZT, Gelber et al.; 2026), whereas latent cell-cycle position may be anchored to a reference trajectory. A common rotation preserves angular differences and concentration. Absolute directions are therefore comparable only when the angular origin and orientation are defined consistently or have been aligned across datasets.

An angle can be represented intrinsically by *θ_i_ ∈* [0, 2*π*) or extrinsically by

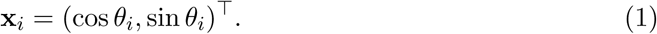

The intrinsic form retains direct angular interpretation, whereas the extrinsic form embeds the angle on the unit circle and enables vector-based analysis while preserving the wraparound property, whereby the endpoints of the angular scale represent the same direction and nearby values across the boundary remain close. Figure 1a illustrates the two representations using angles that are close on the circle but lie on opposite sides of the 0*/*2*π* boundary. For axial data, where opposite directions are equivalent, we use 2*θ_i_* mod 2*π*.

**Figure 1:**
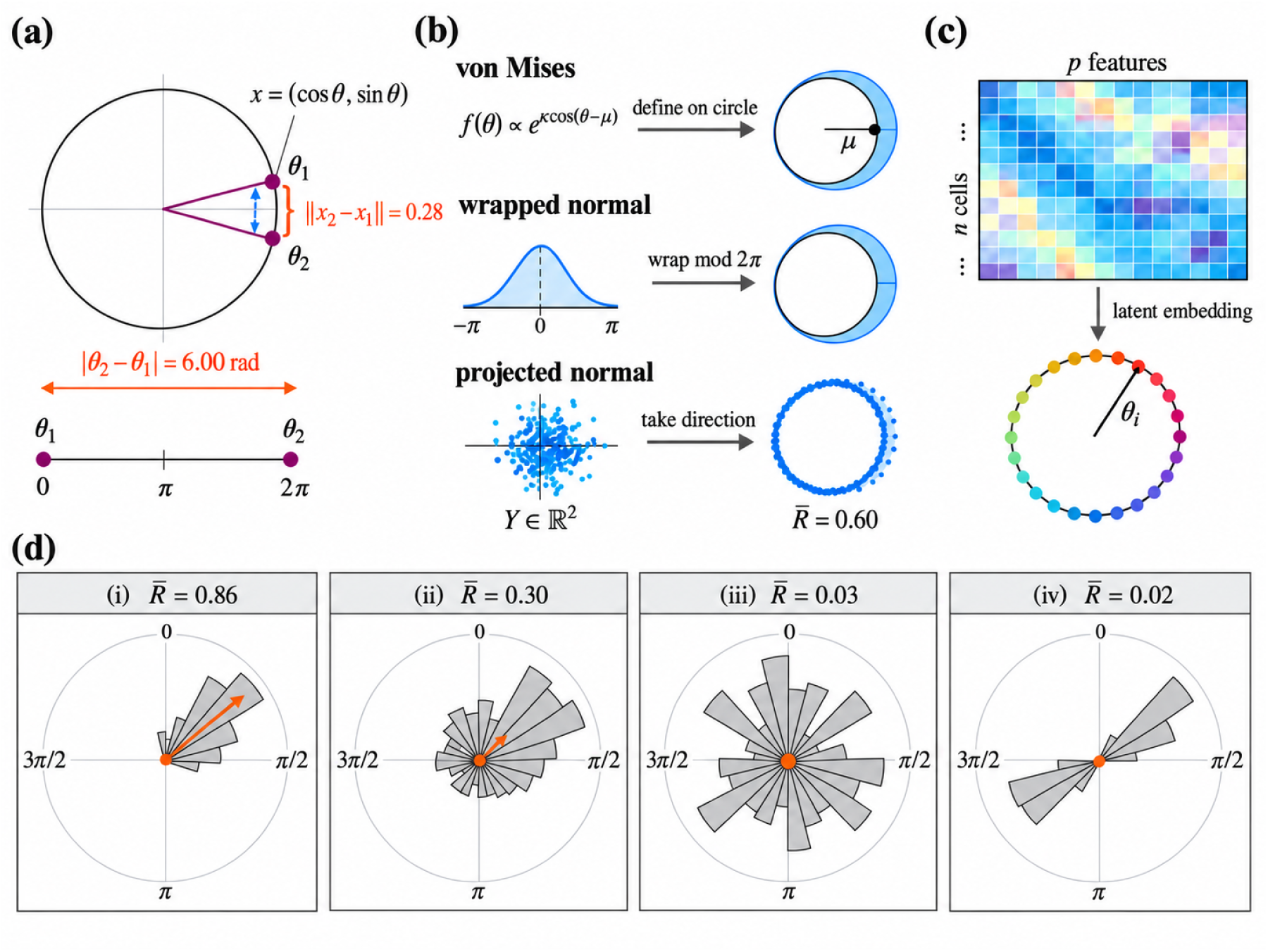
Representing a circular quantity, on simulated data. In (a), two angles lie on opposite sides of the 0*/*2*π* boundary. Their ordinary numerical difference on the linear interval [0, 2*π*) is *|θ*_2_ *− θ*_1_*|* = 6.00 rad, whereas their shortest circular (or geodesic) distance is 2*π − |θ*_2_ *− θ*_1_*|* = 0.28 rad. Under the extrinsic representation in (1), the corresponding Euclidean chord distance is 2 sin(0.28*/*2) *≈* 0.28, illustrating how circular representation preserves proximity across the wraparound boundary. Panel (b) builds a density on the circle in three ways. The von Mises, wrapped, and projected forms are matched at 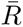 = 0.60, so they differ primarily in distributional shape. Panel (c) infers an angle from an *n × p* profile. In (d), four samples give a feel for the MRL. Arrow length is proportional to 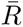, and (iii) and (iv) end up similarly small for different reasons.

More broadly, circular analysis may involve geometric representations, circular probability models, or latent angular variables (Table 2). Figure 1 summarizes these categories, with panel (a) showing intrinsic and extrinsic representations, panel (b) circular probability models, and panel (c) the inference of a latent angle from a cyclic embedding of high-dimensional molecular data. The latter is illustrated by the ovarian-cancer application, where the inferred cell-cycle position is used as the circular variable in downstream analyses (Section 5.2). Figure 1b contrasts von Mises, wrapped, and projected constructions, defined directly on the circle, by wrapping a linear variable modulo 2*π*, or by projecting a

Euclidean vector, respectively. These forms are developed further in Sections 2.4, 3.2, and 4.2. Panel (d) previews the summaries of Section 2.2, and Table 1 maps each analytical objective to a method.

**Table 1:** Practical navigation from an analysis question to the corresponding method, section, and representative R software.

| Question | Analysis | Section | R software |
| --- | --- | --- | --- |
| Is the distribution non-uniform or different across groups? | Uniformity testing and circular group comparison | 2.3 | `circular`,<br>`sphunif` |
| Are there recurrent angular modes? | Finite mixtures of von Mises distributions | 2.4 | `BAMBI` |
| How does a non-circular outcome vary with phase? | Sine-cosine or harmonic regression | 3.1 | base R,<br>`cosinor` |
| How does a circular response vary with covariates? | von Mises or projected-normal regression | 3.2 | `circular`,<br>`brms`, `bpnreg` |
| Is nonlinear prediction the goal? | Tree, forest, or neural-network models with circular encoding | 3.3 | `rpart`, `ranger`,<br>`nnet` |
| Does residual spatial dependence remain after tissue context is considered? | Context-aware circular semivariogram followed, when warranted, by spatial circular modeling | 4 | `CircSpaceTime` |

**Table 2:**
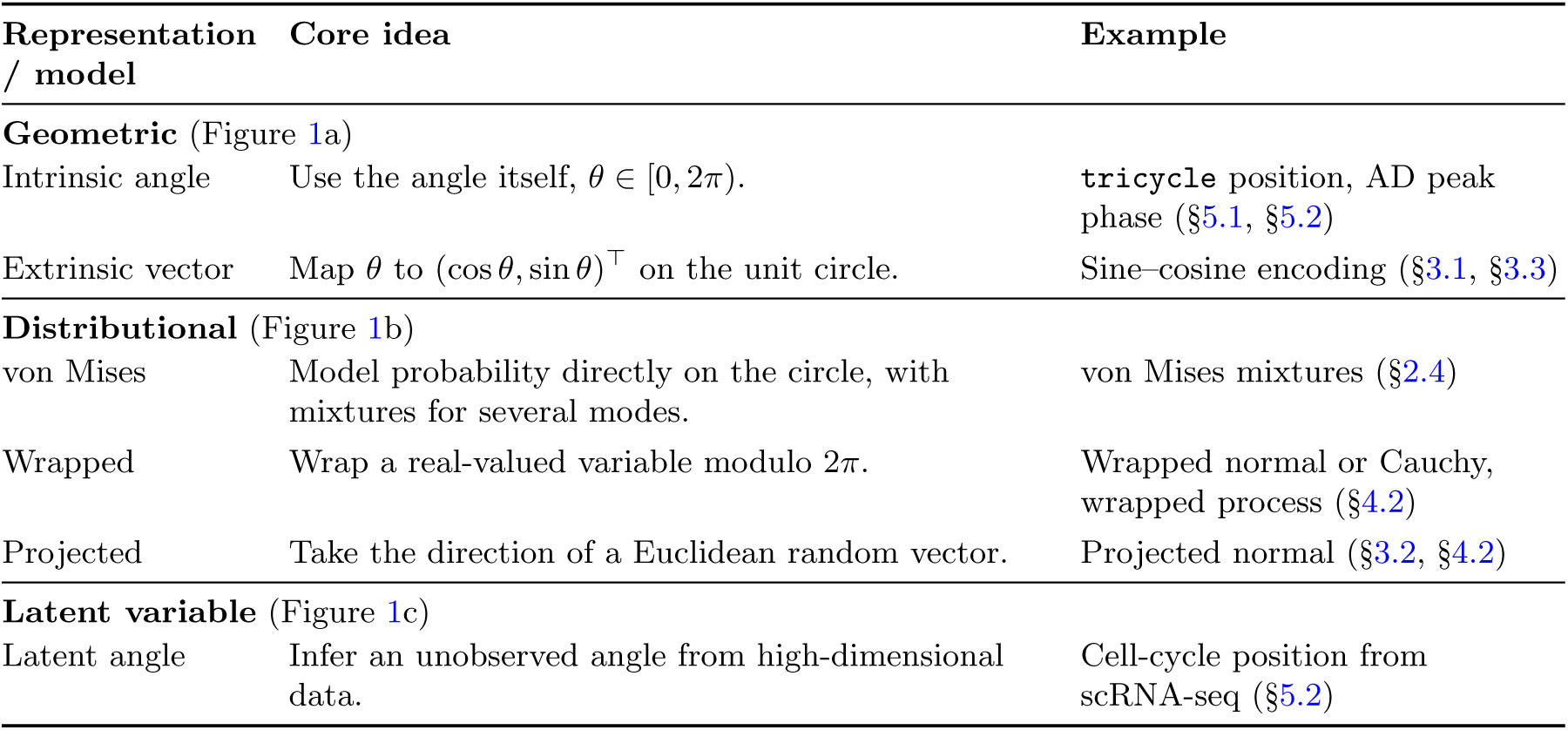
Common representations and probability models for circular quantities. Section numbers point to where each example is developed.

### 2.2 Descriptive statistics and visualization

We summarize a circular distribution using the sample mean direction (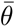) and mean resultant length (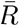, MRL) as follows:

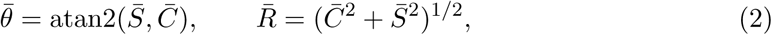

where 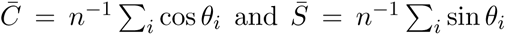, and atan2(*y, x*) denotes the quadrant-aware inverse tangent that returns the direction of the vector (*x, y*). The mean direction describes the average orientation (modulo 2*π*), whereas 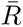*∈* [0, 1] measures first-moment concentration, with larger values indicating stronger directional concentration (Lee; 2010; Pewsey and García-Portugués; 2021). The mean direction is undefined when 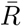 = 0. These quantities are available in the circular package (Agostinelli and Lund; 2025).

We recommend reporting the mean direction and MRL together. A small MRL can represent either a diffuse distribution or cancellation among multiple angular modes. Figure 1d illustrates this distinction. Panels (i) and (ii) show progressively weaker concentration as 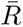 decreases, whereas panels (iii) and (iv) both have near-zero MRL despite representing a diffuse distribution and two opposing modes, respectively. Rose diagrams and circular density plots are therefore useful complements to numerical summaries.

### 2.3 Statistical inference and group comparison

Uniformity testing evaluates whether a circular distribution departs from a uniform distribution. The Rayleigh statistic (Mardia and Jupp; 2000) is based on the first trigonometric moment, 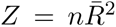, and is well suited to unimodal von Mises-like alternatives (García-Portugués and Verdebout; 2018). The Rayleigh test is implemented in the circular pack-age. However, multimodal distributions can have a small resultant even when they are strongly non-uniform, as in sample (iv) of Figure 1d. Procedures such as the Hermans– Rasson test (Hermans and Rasson; 1985), Pycke test (Pycke; 2010), Kuiper test (Kuiper; 1960), and Watson-type tests (Watson; 1961) provide broader alternatives in this setting. Representative implementations are available in circular and sphunif (García-Portugués and Verdebout; 2026) packages.

Group comparison addresses a different statistical target. Watson-type procedures and design-respecting permutation can compare broader circular distributions. Watson– Williams testing targets differences in mean direction under stronger distributional assumptions and is not a general test of distributional equality (Landler et al.; 2021). MANOVA applied to the sine–cosine representation in (1) provides an alternative when covariates or interactions are required (Landler et al.; 2022). Thus, the inferential target should be specified before testing, whether it concerns mean direction, concentration, or a broader distributional difference.

### 2.4 Circular clustering

Circular clustering identifies recurrent angular modes without predefined group labels. Ordinary clustering of raw angles can separate observations that lie on opposite sides of the wraparound boundary even when they belong to the same mode, for the reason illustrated in Figure 1a. We therefore consider finite mixtures of von Mises distributions,

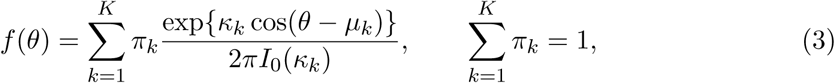

whose kernel is the von Mises density of Figure 1b and where *π_k_*, *µ_k_*, and *κ_k_* denote the component weight, mean direction, and concentration, respectively (Mulder et al.; 2020). The BAMBI package (Chakraborty and Wong; 2021) provides Bayesian mixture estimation and posterior component allocation. Candidate numbers of components can be compared using criteria such as the Watanabe–Akaike information criterion (WAIC) (Watanabe; 2010).

## 3 Regression and prediction with circular variables

Once a circular quantity is defined, the next consideration is whether it serves as a predictor or a response, and whether the goal is association or prediction. Figure 2 summarizes these settings, which are illustrated in the AD and ovarian-cancer applications (Sections 5.1 and 5.2).

**Figure 2:**
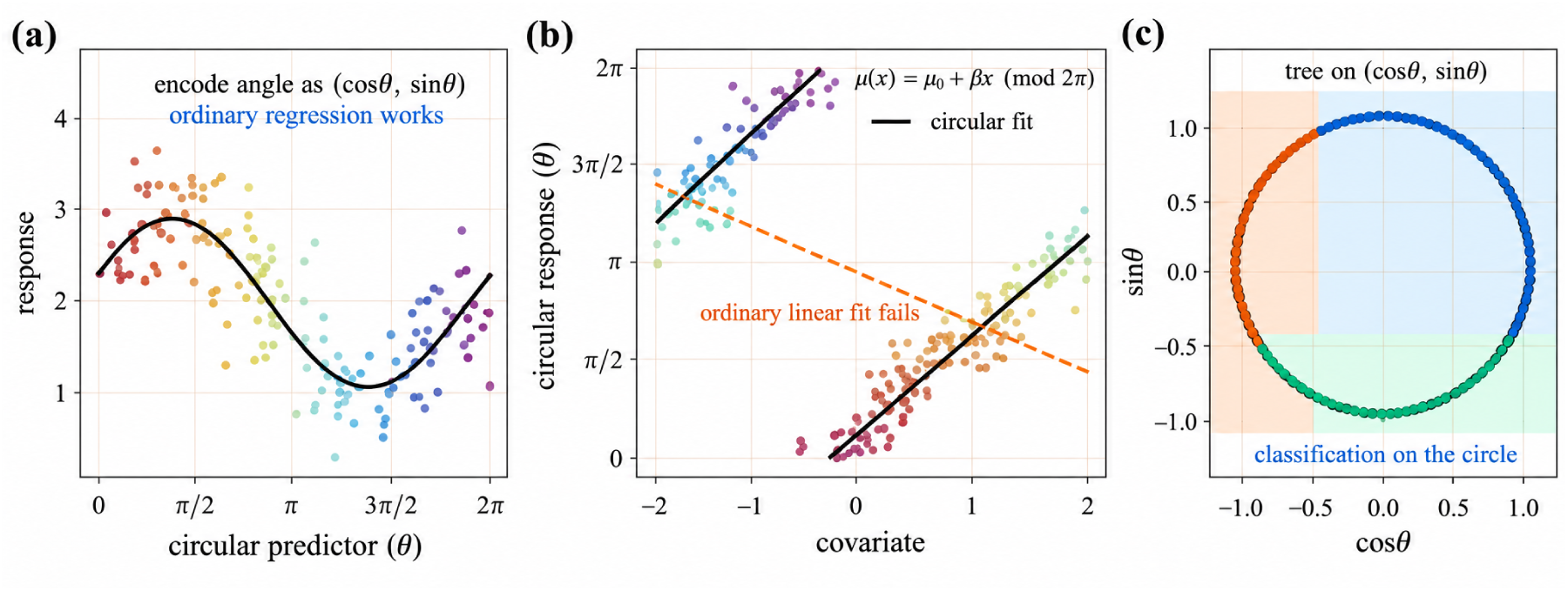
The role of the angle decides the model, on simulated data. Panel (a) takes the angle as a predictor, handled by the harmonic model of (4). In (b), the angle is the response, handled by von Mises regression. Its mean direction wraps, so the least-squares line (dashed) leaves the data where the circular fit (solid) follows it. Panel (c) shows a classification tree using sine–cosine encoding of the circular predictor, where axis-aligned splits correspond to arcs on the circle.

### 3.1 Circular predictors: harmonic regression

When *θ_i_* is a predictor and *Y_i_* is non-circular, we represent the circular effect through its sine and cosine components. A first-order harmonic model is

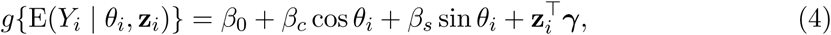

where **z***_i_* contains additional covariates and ***γ*** is the corresponding coefficient vector. The two circular coefficients, *β_c_*and *β_s_*, should generally be interpreted and tested jointly. Figure 2a shows the fitted model for a simulated non-circular response. Higher harmonics can be added when the relationship has a more complex periodic form (Yang and Su; 2010). Standard linear and generalized linear models can be fitted using lm and glm, whereas cosinor regression provides an equivalent representation of periodic effects in terms of amplitude and acrophase (Nelson et al.; 1979; Sachs; 2023).

### 3.2 Circular responses: von Mises and projected-normal regression

When *θ_i_* is the response, ordinary linear regression is inappropriate because fitted values do not respect the circular boundary. Figure 2b shows the consequence when the conditional mean direction wraps across the covariate range. One approach is von Mises regression, which models a predictor-dependent mean direction (Fisher and Lee; 1992). Frequentist estimation is available through lm.circular() in the circular package. Bayesian von Mises regression can be fitted with brms (Bürkner; 2017; Carpenter et al.; 2017). These models are most suitable when the conditional circular distribution is approximately symmetric and unimodal.

Projected-normal regression provides an alternative in which covariates can shift both the location and the concentration of the angle. It makes the projected construction of Figure 1b depend on covariates by defining a latent bivariate normal response and taking the observed angle as its direction,

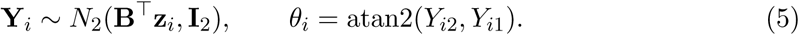

Here, **Y***_i_* = (*Y_i_*_1_*, Y_i_*_2_)*^⊤^* is a latent bivariate variable, **z***_i_* is the predictor vector, and **B** is the regression coefficient matrix. The covariance is fixed at the identity so that the model is identifiable. The model is implemented in the bpnreg package (Cremers; 2024). Because covariates act on the latent vector rather than on the angle itself, coefficients should not be interpreted as ordinary linear slopes. We instead translate fitted effects back to the circular scale (Cremers et al.; 2018). For categorical predictors, wrapped differences in fitted mean direction can be reported relative to a reference level and, for circadian data, converted to hours.

### 3.3 Machine learning: circular encoding and validation

Machine-learning methods provide flexible alternatives when nonlinear effects or interactions are important, particularly when prediction is the primary goal, although their use is not limited to predictive tasks. When the circular quantity is a predictor, (1) allows standard learners to use it directly. Decision trees implemented in rpart (Therneau and Atkinson; 2025) provide interpretable partitions of the sine–cosine plane, corresponding to contiguous arcs of the circle (Figure 2c), whereas random forests implemented in ranger (Wright and Ziegler; 2017) aggregate many trees and provide permutation-based variable importance (Breiman; 2001; Cutler et al.; 2007; Wright and Ziegler; 2017). These approaches are illustrated in the ovarian-cancer and AD applications, respectively (Sections 5.2 and 5.1).

When the circular quantity is the response, its sine and cosine components can instead be predicted and combined as 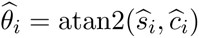. This approach avoids the 0*/*2*π* discontinuity and can be implemented with a neural network such as nnet (Venables and Ripley; 2002), as illustrated for AD peak-phase prediction in Section 5.1. Held-out performance can then be evaluated with angular error, chord loss, or circular correlation. Squared-error prediction of the two components, however, can favor the dominant resultant direction when the conditional distribution is broad or multimodal.

## 4 Spatial circular analysis

Spatial omics allow circular quantities to be examined directly on tissue, provided that the angle and spatial coordinate are defined on the same observational unit, such as a cell or spot (Section 2.1). A phase estimated after aggregation across spots, for example, should not be mapped back to individual locations. Even at matched resolution, spatially similar angles do not necessarily imply residual spatial dependence because tissue regions can themselves induce spatial structure.

### 4.1 Composition before covariance

Tissues contain biologically distinct regions, such as tumor, stroma, or anatomical layers, that may have different angular distributions. Let *G_i_* denote the tissue region of observation *i*. Under a composition-only mechanism,

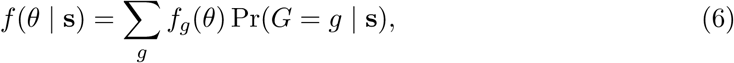

where *f_g_*is the angular distribution in region *g*. The marginal distribution can therefore vary across tissue simply because region membership changes with location, even when observations are independent within each region. Figure 3a illustrates this effect. Angles are sampled independently from the region-specific distributions in Figure 3c, yet the map appears spatially organized because the regions themselves are organized. In contrast, Figure 3b shows residual spatial dependence within a single region, where nearby observations remain similar as the local mean direction changes smoothly across space.

**Figure 3:**
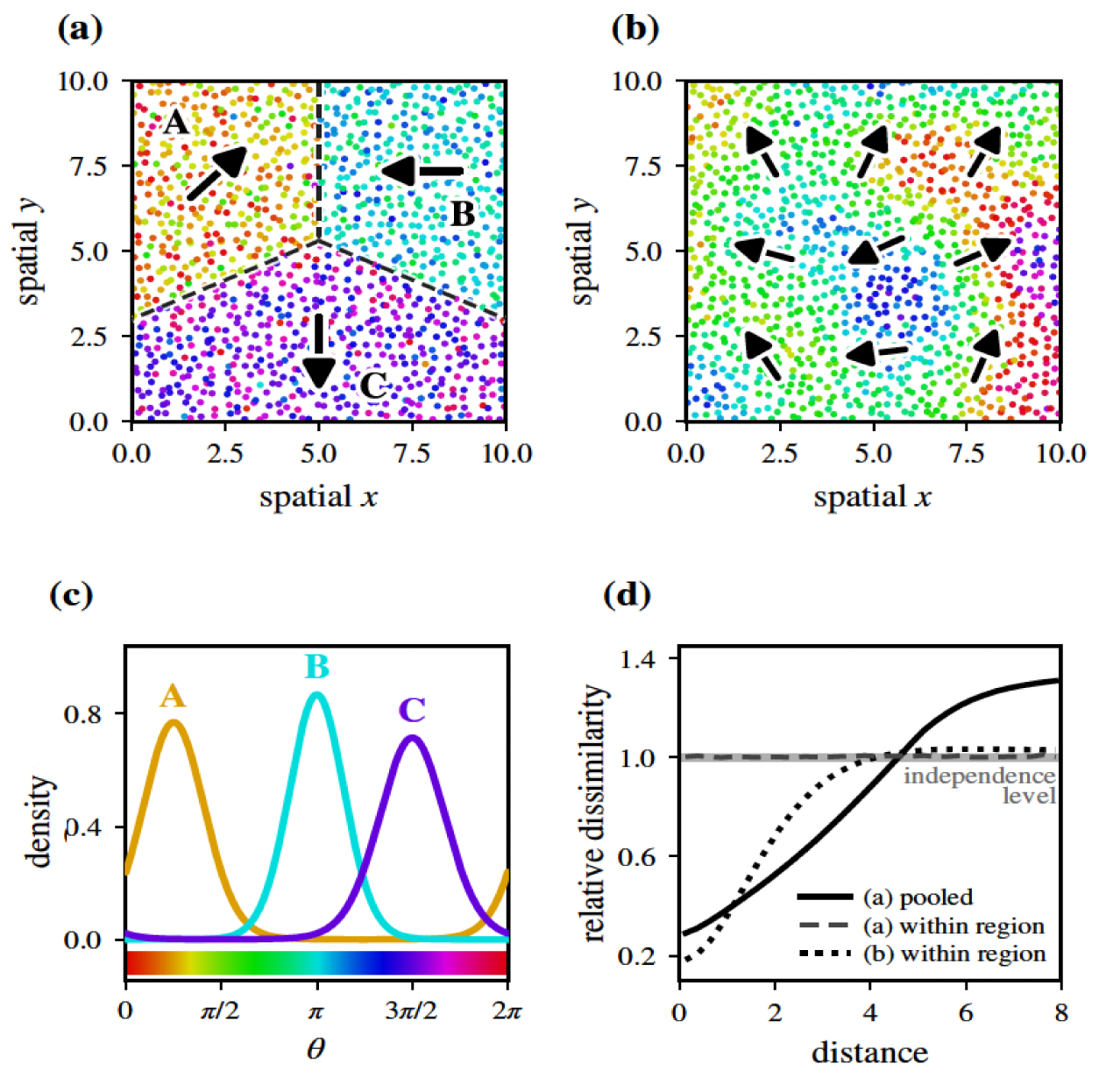
Composition before covariance, on simulated tissue. (a) Three tissue regions (A, B, and C, separated by dashed lines) with angles drawn independently from region-specific distributions. (b) A single region with a spatially varying mean direction. Arrows indicate region-specific mean directions in (a) and local mean directions in (b). (c) Angular distributions for the three regions in (a). The color bar shows the cyclic scale used in (a) and (b). (d) Relative dissimilarity as a function of distance. A value of one indicates independence. The apparent short-range similarity in (a) disappears within regions, whereas within-region dependence persists in (b).

This distinction motivates an ordering that we call composition before covariance. Angular distributions are first compared between tissue regions, and spatial covariance is modeled only when local similarity remains within a region. This ordering avoids attributing region-driven structure to residual spatial dependence.

To assess local similarity, we measure the angular dissimilarity of observations *i* and *j* by

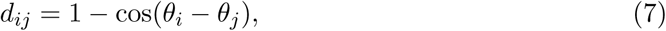

which ranges from zero for identical directions to two for opposite directions. Averaging *d_ij_* over pairs at similar spatial distances yields an empirical circular semivariogram. Because *d_ij_* depends only on angular differences, it is invariant to a common rotation of the reference system. For independent observations from the same circular distribution,

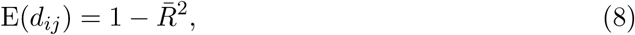

where 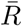 is the mean resultant length (Section 2.2). Dividing the empirical semivariogram by this independence level gives the relative dissimilarity shown in Figure 3d, for which a value of one corresponds to independence. Values below one at short distances indicate excess local angular similarity.

In practice, the diagnostic proceeds in four steps. First, map the circular quantity together with tissue regions. Second, compare angular distributions across regions. Third, examine pooled relative dissimilarity. Finally, repeat the calculation within sufficiently represented regions. In Figure 3d, the simulation from panel (a) shows apparent shortrange similarity when pooled but not within regions, whereas the within-region curve from panel (b) remains below one at short distances and approaches one with distance. Thus, disappearance of short-range similarity within regions supports composition, whereas persistence supports residual spatial dependence and motivates the models of Section 4.2.

### 4.2 Spatial circular models and prediction

When residual spatial dependence remains, wrapped and projected Gaussian processes provide established models for spatial circular data (Jona-Lasinio et al.; 2012; Wang and Gelfand; 2014). A wrapped Gaussian process maps a real-valued spatial process onto the circle, whereas a projected Gaussian process defines the angle from a latent bivariate process,

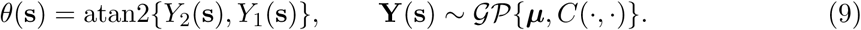

Here, **Y**(**s**) = (*Y*_1_(**s**)*, Y*_2_(**s**))*^⊤^*, ***µ*** is its mean vector, and *C*(**s**, **s***^′^*) describes spatial dependence. These models extend the wrapped and projected distributions in Figure 1b to spatial data, with (9) corresponding to the spatial extension of (5).

The CircSpaceTime package implements projected-process fitting with ProjSp() and prediction with ProjKrigSp() (Jona Lasinio et al.; 2020). Prediction should be evaluated at spatially separated held-out locations using circular metrics such as angular error or circular continuous ranked probability score (Grimit et al.; 2006).

## 5 Case studies

We now apply the workflow to two spatial transcriptomic datasets whose circular quantities differ in provenance and observational unit (Section 2.1 and Table 2). The AD example uses a derived gene-by-context peak phase and follows Sections 2 and 3. The ovarian-cancer example uses a latent cell-level position and also follows Section 4. Table 1 lists the method and R package for each analysis, and Supplementary Table S1 links each analysis to its supplementary results and code.

### 5.1 Derived circadian peak phase in AD

#### Data and circular quantity

The AD analysis uses 65 mouse-brain spatial transcriptomic samples spanning WT and APP23 genotypes, 7- and 14-month ages, both sexes, and multiple Zeitgeber times (ZT). Spots were assigned to brain regions and aggregated to pseudobulk counts for each sample and region. Within each brain region, genotype, and age stratum, hereafter a context, a log-link form of the harmonic model in (4) was fitted for each gene with DESeq2 (Love et al.; 2014),

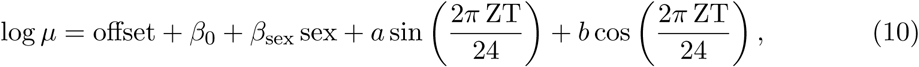

where *µ* is the expected pseudobulk count. Rhythmicity was tested with a likelihood ratio test against a reduced model without the harmonic terms, retaining 16,390 gene-by-context estimates with adjusted *p <* 0.10 (Supplementary Section S3.1). For gene *g* in context *c*, rhythmic amplitude and peak phase (acrophase) were then obtained from the two harmonic coefficients (Section 3.1),

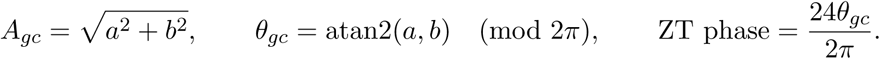

Because DESeq2 reports *a* and *b* on the log2 scale, *A_gc_* is a log2 amplitude. The circular quantity is thus a derived peak phase anchored to ZT0 and defined for gene-by-context estimates rather than spots (Section 2.1), so the spatial analyses of Section 4 do not apply. Figure 4 shows representative rhythmic genes in cortex layer 2/3 at 7 months, and Supplementary Figure S1 shows subregional heterogeneity in peak phase.

**Figure 4:**
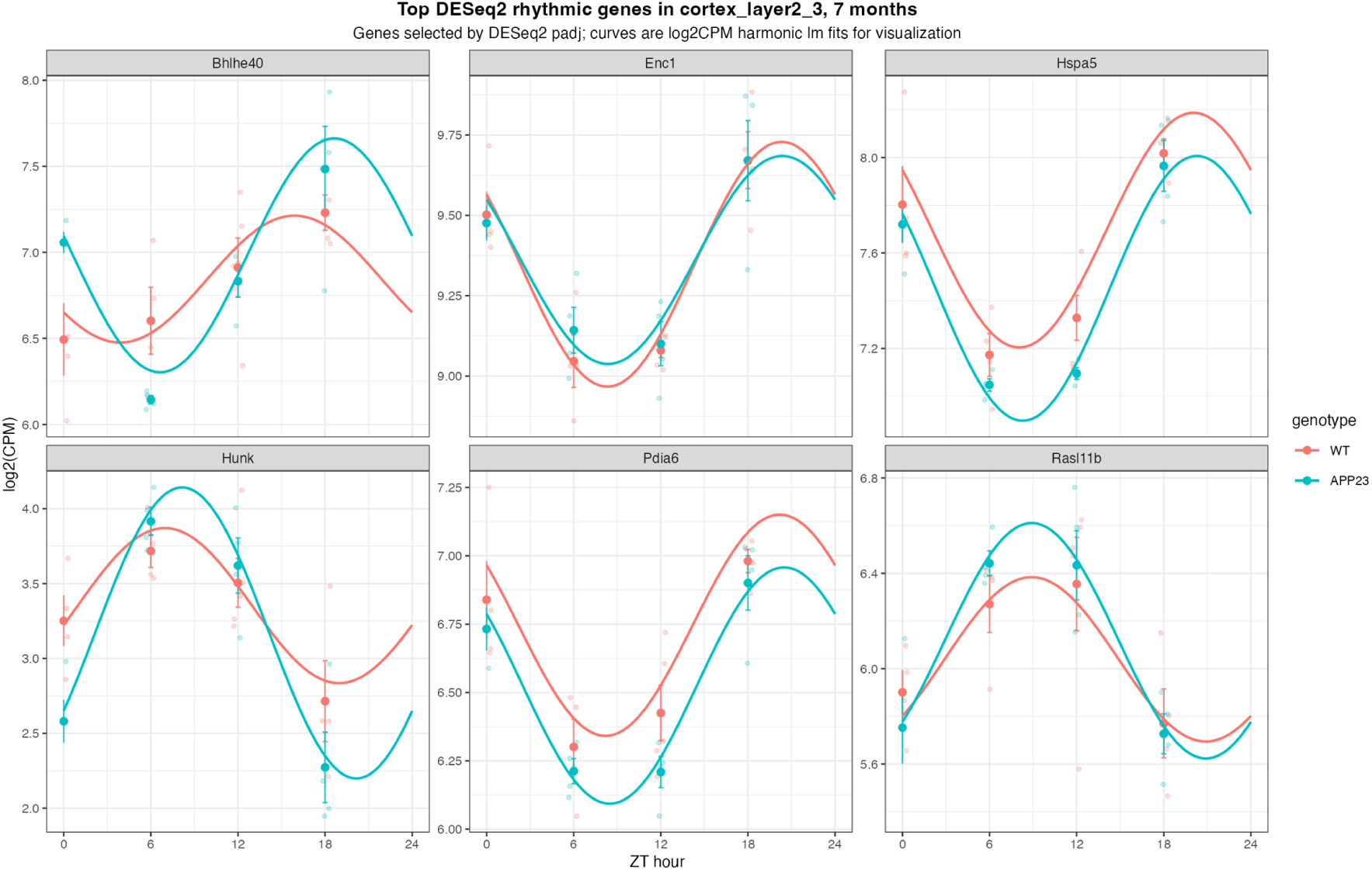
Derivation of circadian peak phase in the AD example. Representative rhythmic genes in cortex layer 2/3 at 7 months illustrate genotype-specific harmonic expression patterns. Peak phase and rhythmic amplitude of each gene in each context were derived from the fitted sine and cosine coefficients of the harmonic model in (10).

#### Description and circular clustering

Peak-phase distributions were broad and partly multimodal across brain regions, genotypes, and ages (Figure 5a). The pooled MRL in (2) was only 0.10, reflecting partial cancellation between a dominant late dark-phase mode and a weaker, roughly antiphase mode, as in sample (iv) of Figure 1d. A single mean direction is therefore an incomplete summary (Section 2.2). Phase composition also varied across contexts (Figure 5b).

**Figure 5:**
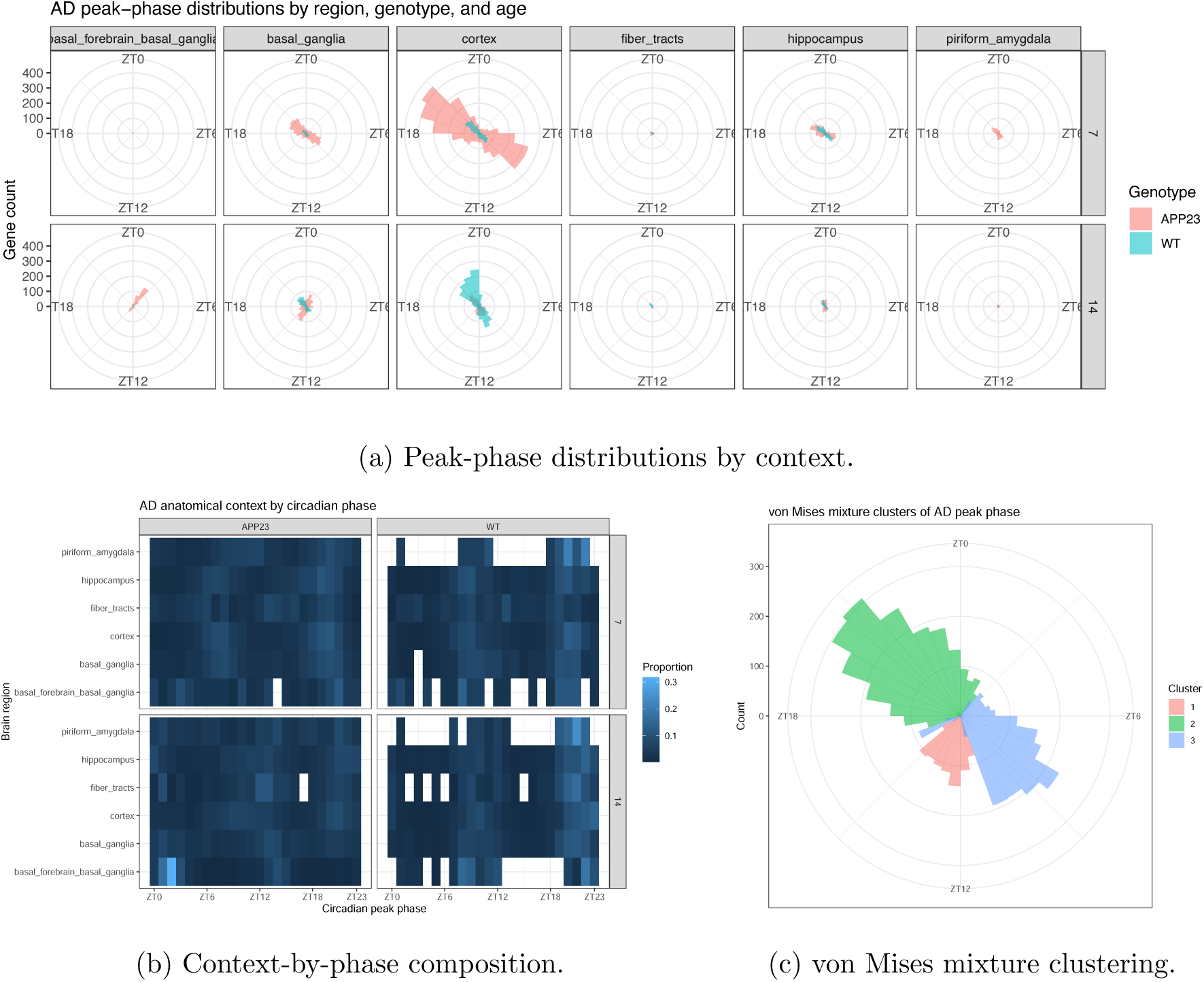
Distributional structure of derived circadian peak phase in the AD example. (a) Rose diagrams show peak-phase distributions across brain regions, genotypes, and ages. (b) The context-by-phase heatmap summarizes differences in relative phase composition across contexts. (c) A three-component von Mises mixture represents recurrent modes in the pooled peak-phase distribution. Together, these panels show broad multimodal structure that a single mean direction does not adequately summarize.

We fitted the von Mises mixture in (3) with BAMBI (Section 2.4) to 5,000 randomly sampled peak phases and compared one to four components by WAIC. The selected three-component solution had mean directions near ZT13.2, ZT21.1, and ZT8.1, accounting for about 16%, 48%, and 35% of the sampled phases (Figure 5c), with similar proportions across the six largest brain regions (Supplementary Table S3). The two major modes lie in the late dark phase (ZT21) and the mid-light phase (ZT8), an antiphase pair resembling the dawn- and dusk-anticipatory peaks reported across mouse tissues (Zhang et al.; 2014), and their regional stability indicates a brain-wide temporal architecture. The dominant ZT21 mode was less prevalent in APP23 than in WT (45% vs 55%, Supplementary Table S3).

#### Peak phase as a circular predictor

We first used peak phase as a circular predictor of rhythmic amplitude, taking *Y_gc_* = *A_gc_* as the non-circular response and genotype, age, and brain region as the additional covariates **z***_c_*. With an identity link, the harmonic model (4) becomes

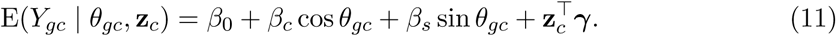

A joint *F*-test of *β_c_* and *β_s_* (Section 3.1) showed a strong association between peak phase and amplitude after adjustment for context (*F* = 309.69, *p <* 2.2 *×* 10*^−^*^16^).

A random forest fitted with ranger (Section 3.3) then predicted *Y_gc_* from sin *θ_gc_*, cos *θ_gc_*, and **z***_c_*. The circular features had the highest permutation importance (Supplementary Figure S2), but held-out performance was modest (correlation 0.220, RMSE 0.130).

#### Peak phase as a circular response

We next modeled peak phase as the response (Section 3.2), using the gene-by-context estimates with *A_gc_* above its median and z-scoring amplitude across these estimates to obtain Ã*_gc_*. Von Mises regression with circular::lm.circular() (Fisher and Lee; 1992) failed increasingly often as the sample size grew (Supplementary Table S4), so it served only as a benchmark on subsamples. Bayesian von Mises regression with brms (Bürkner; 2017) and projected-normal regression with bpnreg::bpnr() (Cremers; 2024) used Ã*_gc_* and **z***_c_* as predictors, which for the projected-normal model gives

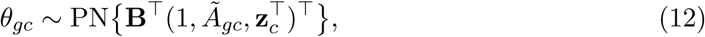

where PN denotes the projected normal distribution of (5), that is, the direction of a bivariate normal vector with the given mean and identity covariance.

Because the brms chains did not converge (Supplementary Figure S3a), the projected-normal model, which converged and improved on its intercept-only fit (Supplementary Figure S3b, Supplementary Table S5), was retained. Its effects are reported as phase shifts in hours relative to the reference condition, with 95% highest posterior density (HPD) intervals (Figure 6a). The dominant peak was about 1.3 h earlier in APP23 than in WT (95% HPD *−*1.5 to *−*1.1 h), consistent with the circadian disruption reported for this model (Gelber et al.; 2026), but about 0.6 h later at 14 than at 7 months, so AD pathology and aging shift peak timing in opposite directions. Regional differences stayed within about 1 h. Because peak phase is multimodal (Figure 5c), these shifts describe the dominant timing trend rather than a shift of every rhythmic gene.

**Figure 6:**
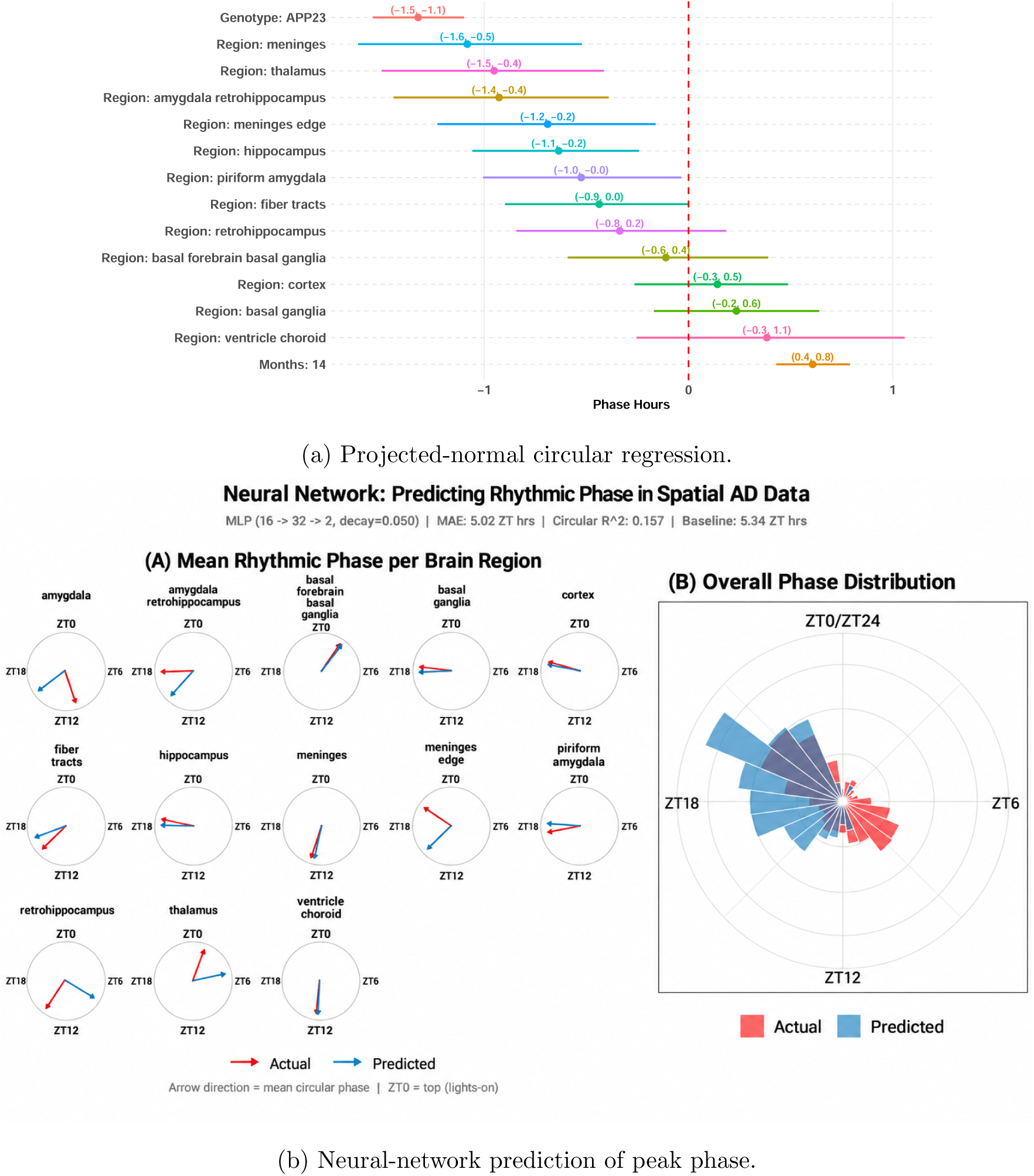
Model-based analysis of derived circadian peak phase as a circular response in the AD example. (a) Projected-normal regression summarizes phase shifts in hours for genotype, age, and brain region relative to the reference condition, with 95% HPD intervals. (b) Neural-network prediction compares observed and predicted peak phase in the held-out test set, showing regional mean directions and the corresponding overall phase distributions. The neural network captures the dominant phase structure but only modestly improves prediction over the mean-direction baseline.

#### Neural-network prediction of peak phase

A multilayer perceptron fitted with nnet (Section 3.3) predicted (sin *θ_gc_,* cos *θ_gc_*) from *A_gc_* and **z***_c_*, and peak phase was recovered with atan2. After dummy encoding, the network had 16 inputs, one hidden layer of 32 units, and two linear outputs. The gene-by-context estimates were split into training, validation, and test sets (60%, 20%, and 20%), and weight decay was selected on the validation set from *{*0.001, 0.01, 0.05*}*, giving 0.05.

On the test set, the mean absolute angular error was 5.02 h, compared with 5.34 h for a baseline that predicts the training-set mean direction (2) of each brain region and genotype group, and the circular *R*^2^ based on chord loss was 0.157. The network thus improved only modestly on the baseline. The predicted distribution captured the dominant mode but smoothed the weaker secondary mode (Figure 6b), as expected for squared-error prediction of sine and cosine components under a multimodal distribution (Section 3.3). This is consistent with substantial gene-level heterogeneity beyond the measured context.

### 5.2 Latent cell-cycle position in ovarian cancer

#### Data and circular quantity

The second example uses the public 10x Genomics Xenium Prime FFPE human ovarian-cancer dataset (10x Genomics; 2024). Each annotated cell *i* has tissue coordinates **s***_i_*, a pathology region *G_i_* (Section 4.1), a discrete Seurat cell-cycle phase (G1, S, G2M, or undecided), and a continuous cell-cycle position inferred by tricycle (Zheng et al.; 2022). tricycle projects each expression profile onto a pretrained two-dimensional embedding (*u_i_*_1_*, u_i_*_2_) and takes its direction as the angle,

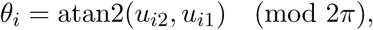

the latent-angle construction of Figure 1c. In this reference frame, positions near 0 or 2*π* correspond approximately to G0/G1, 0.5*π* to G1/S, and *π* to S/G2–M. Because each angle is paired with cell coordinates, the observational-unit condition of Section 4 holds.

#### Description, uniformity testing, and circular clustering

Figure 7 maps pathology regions, cell-cycle position on a cyclic color scale (Section 2.2), cell type, and Seurat phase. The continuous position varies throughout the tissue, whereas G1 is broadly distributed and S and G2M occupy more localized subsets. Across all 113,295 annotated cells, the mean direction and MRL in (2) were 0.27*π* and 0.151. Tumor cells had a broad, weakly concentrated distribution. Blood vessels, immune cells, and fallopian-tube cells were concentrated near G0/G1, and the small necrosis region lay near S/G2 (Table 3A and Supplementary Figure S4a). A high MRL means cells share a similar cell-cycle position, whereas a low MRL means they are spread across the cycle. Blood vessels and immune cells therefore appear mostly resting in G0/G1, whereas tumor cells are dividing at many different stages at once. The S/G2 position of necrotic cells is likely an artifact of degraded RNA rather than true cell division.

**Figure 7:**
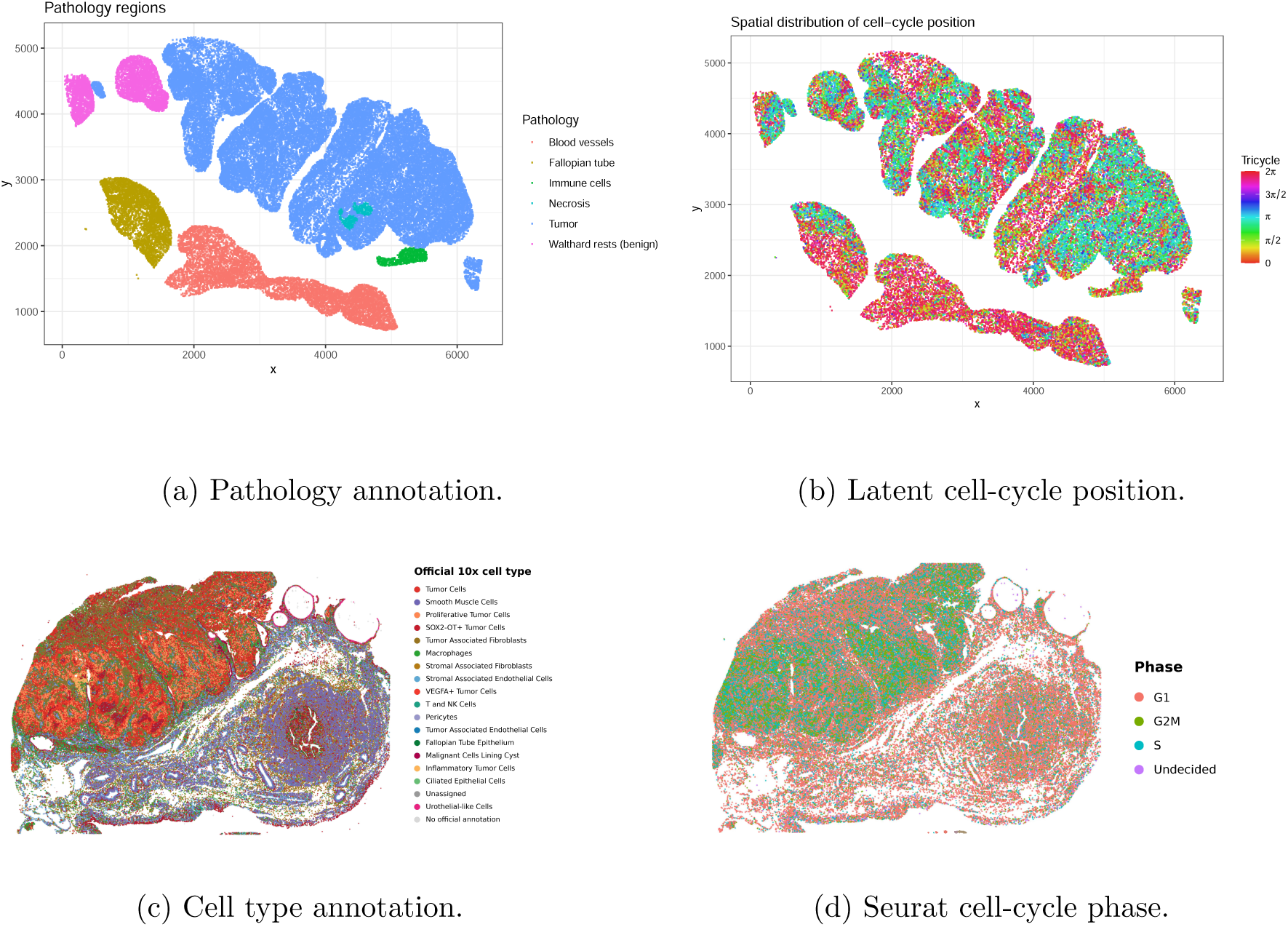
Tissue context and cell-cycle state in the ovarian-cancer example. (a) Spatial distribution of pathology annotations across the Xenium specimen. (b) Cell-level spatial distribution of continuous tricycle cell-cycle position. (c) Cell-type annotations provide complementary tissue context. (d) Discrete Seurat cell-cycle classifications show the corresponding G1, S, G2M, and undecided states. The continuous tricycle position and discrete Seurat phase provide complementary representations of the same transcriptional cell-cycle program.

**Table 3:** Circular summaries of tricycle cell-cycle position in the ovarian-cancer example. (A) Mean direction, mean resultant length (MRL), and Rayleigh statistic 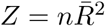 for all annotated cells by pathology region. (B) MRL and Rayleigh *Z* across two subsamples and the full data for tumor cells and Walthard rests, illustrating the dependence of the Rayleigh statistic on observational sample size despite similar concentration.

| **(A) Circular summaries by pathology region** |  |  |  |  |
| --- | --- | --- | --- | --- |
| **Pathology** | **$n$** | **Mean direction** | **MRL** | **Rayleigh $Z$** |
| Tumor | 79,104 | $0.51\pi$ (G1/S) | 0.133 | 1,408 |
| Blood vessels | 15,558 | $0.01\pi$ (G0/G1) | 0.611 | 5,808 |
| Walthard rests (benign) | 8,479 | $0.60\pi$ (G1/S) | 0.135 | 154 |
| Fallopian tube | 8,014 | $0.09\pi$ (G0/G1) | 0.296 | 703 |
| Immune cells | 1,285 | $0.13\pi$ (G0/G1) | 0.495 | 315 |
| Necrosis | 855 | $0.91\pi$ (S/G2) | 0.479 | 196 |
| All annotated cells | 113,295 | $0.27\pi$ (G0/G1) | 0.151 | 2,597 |

Table 3: Circular summaries of `tricycle` cell-cycle position in the ovarian-cancer example.
| **(B) Effect of observational sample size: MRL / $Z$ ($n$)** |  |  |  |
| --- | --- | --- | --- |
| **Pathology** | **$\sim 1,500$-cell sample** | **5,000-cell sample** | **All annotated cells** |
| Tumor | 0.155 / 19.9 (827) | 0.126 / 54.6 (3,425) | 0.133 / 1,408 (79,104) |
| Walthard rests (benign) | 0.132 / 1.6 (92) | 0.150 / 9.2 (409) | 0.135 / 154 (8,479) |

Uniformity was summarized with the Rayleigh statistic 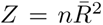 (Section 2.3). Tumor cells and Walthard rests had nearly identical MRLs (0.133 and 0.135) but very different *Z* because their cell numbers differ, and subsampling shows *Z* rising sharply with *n* while the MRL stays stable (Table 3B). The MRL is therefore reported as an effect size alongside significance.

We fitted the von Mises mixture in (3) with BAMBI (Section 2.4) to 5,000 randomly sampled cells and compared one to four components by WAIC. The selected three-component solution was centered near 1.95*π* (G0/G1), 0.58*π* (G1/S), and 1.05*π* (S/G2–M), accounting for about 49%, 24%, and 28% of the sampled cells (Figure 8a). Membership agreed with the Seurat labels, with 79% of G1 cells in component 1, 48% of S cells in component 2, and 77% of G2M cells in component 3 (Supplementary Table S6). Because Seurat phase uses related cell-cycle genes, this agreement supports but does not independently validate the modes. Component prevalence differed across pathology regions (Figure 8b), with blood vessels dominated by component 1 (81%) and tumor cells spread across all three (43%, 26%, and 32%). Read as a cycling fraction, about 57% of tumor cells fall outside the G0/G1 mode versus 19% of blood-vessel cells (Supplementary Table S6), a label-free analogue of a proliferation index.

**Figure 8:**
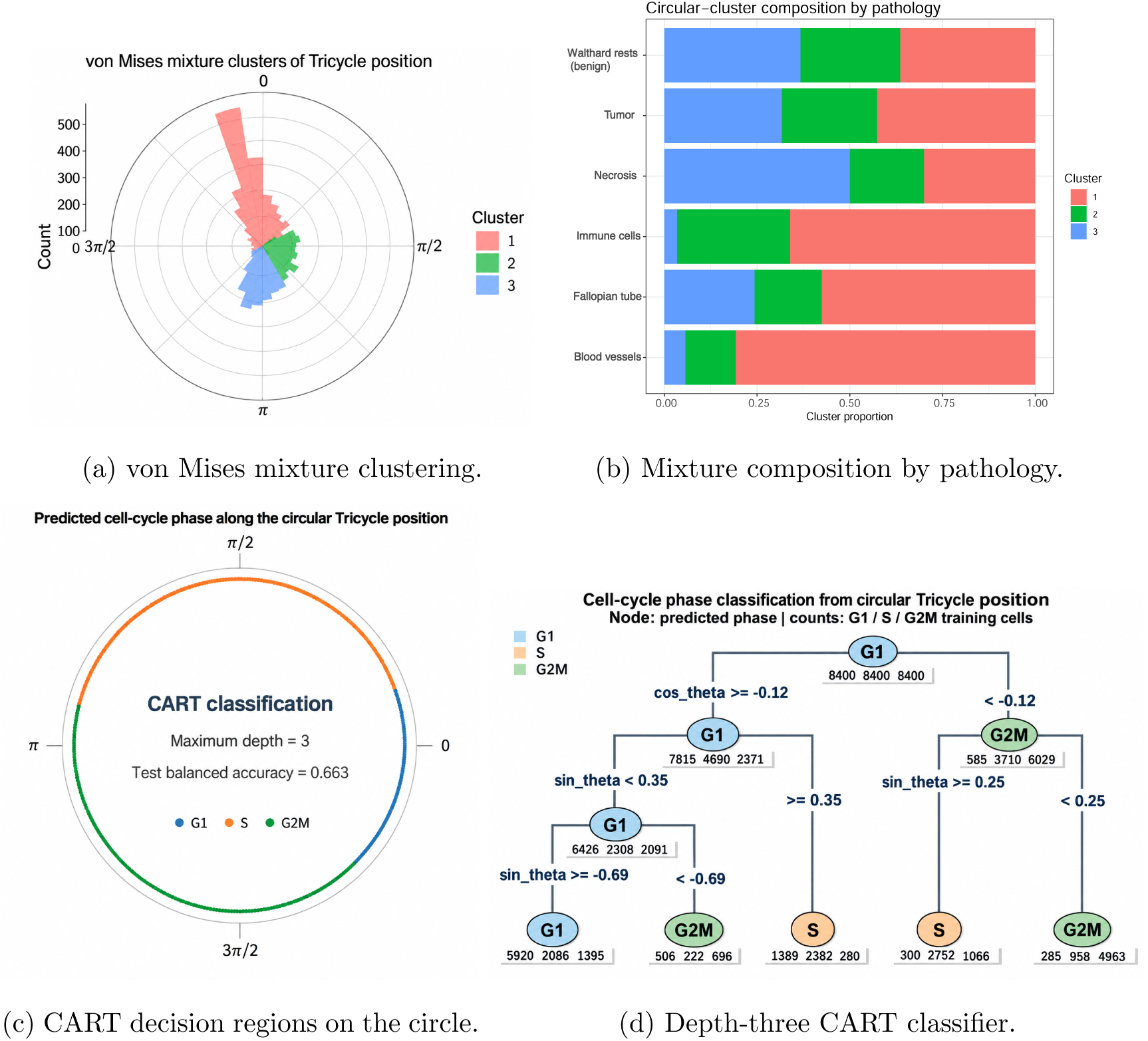
Circular clustering and classification of latent cell-cycle position in the ovarian-cancer example. (a) The selected three-component von Mises mixture identifies recurrent angular modes of tricycle position. (b) The relative prevalence of the three mixture components varies across pathology regions. (c) CART decision regions projected onto the circle show how the sine–cosine representation partitions cell-cycle position into regions associated with discrete Seurat phases. (d) The corresponding depth-three classification tree displays the sequential splits on cos *θ* and sin *θ*.

#### Cell-cycle position as a circular predictor

We next used cell-cycle position to predict Seurat phase (G1, S, or G2M). Following Section 3.3, a CART model fitted with rpart used the extrinsic coordinates (cos *θ_i_,* sin *θ_i_*) in (1) as predictors, with a stratified 70:30 training– test split, a class-balanced training set of 25,200 cells (8,400 per class), and depth limited to three. The first split at cos *θ ≈ −*0.12 and later splits on sin *θ* (Figure 8d) divide the circle into contiguous arcs dominated by G1, S, and G2M (Figure 8c), as in Figure 2c. Held-out balanced accuracy, which accounts for class imbalance, was 0.663.

#### Composition before covariance

Pathology regions differ in cell-cycle position (Table 3A) and occupy distinct parts of the tissue (Figure 7a), so the pattern in Figure 7b could arise from composition alone, as in Figure 3a. Having mapped and compared regions above, we completed the diagnostic of Section 4.1 on about 1,500 cells sampled proportionally across pathology regions (Figure 9a). The empirical circular semivariogram based on (7) was computed for all sampled cells and for tumor cells only and compared with the independence level in (8).

**Figure 9:**
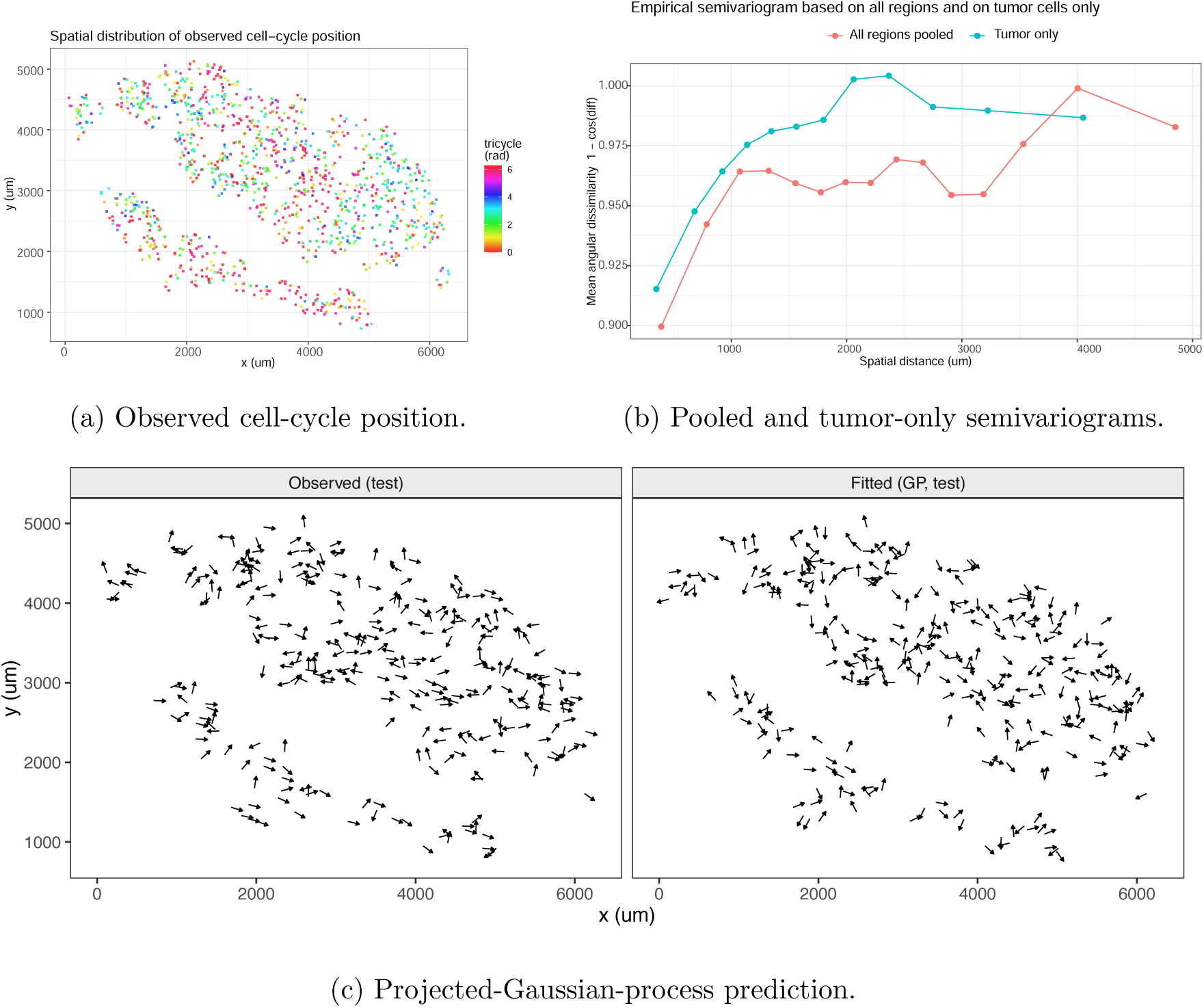
Spatial analysis of latent cell-cycle position in the ovarian-cancer example, following Section 4. (a) Observed tricycle position in the proportional subsample of about 1,500 cells. (b) Empirical circular semivariograms based on (7) for all sampled cells and for tumor cells only. Short-range similarity persists within the tumor region, the residual-dependence outcome of Figure 3d. (c) Observed directions at 300 held-out cells and predictions by projected-Gaussian-process kriging based on (9).

The pooled curve increased with distance, and short-range similarity persisted within tumor cells (Figure 9b). The tumor-only curve stayed below its independence level at short distances and reached it at about 1–2 mm, matching the residual-dependence outcome in Figure 3d. The pattern is therefore not explained by the arrangement of pathology regions alone, and instead means that neighboring tumor cells within about 1–2 mm tend to be at similar cell-cycle stages, so proliferation is patchy within the tumor. The effect is modest and comes from one specimen, and it may reflect local biological coordination, unmeasured cell states, or technical variation, so additional specimens are needed before interpreting it as a reproducible tumor-microenvironment pattern.

#### Spatial prediction of cell-cycle position

We then fitted the projected Gaussian process in (9) (Section 4.2) with a constant mean and exponential correlation using CircSpaceTime::ProjSp() (Jona Lasinio et al.; 2020). The subsample was split into 1,199 training and 300 held-out cells, and held-out angles were predicted by circular kriging with ProjKrigSp(). Four chains were run for 20,000 iterations with 10,000 discarded as burn-in (Supplementary Section S5.2). The kriged directions reproduce the locally dominant cell-cycle position while smoothing cell-level variation (Figure 9c). Randomly held-out cells can lie next to training cells and overstate accuracy, and a single mean and covariance across pathology regions can absorb compositional differences (Section 4.1). Overall, treating cell-cycle position as an angle showed three things that the G1, S, and G2M labels alone could not. Resting and dividing compartments differ in how tightly their cells cluster on the cycle, the share of dividing cells can be estimated without labels, and neighboring tumor cells within about 1–2 mm tend to be at similar stages.

### 5.3 Practical considerations

The two applications motivate four recommendations for circular data analysis in spatial omics. First, match the analysis and display to the observational unit (Section 2.1). Cell-level tricycle positions can be mapped on tissue (Figure 7b), whereas AD peak phases summarize genes within contexts and should be displayed at that level (Figure 5b). Report how *θ_i_* was obtained, its angular origin and orientation, and any filtering or estimation uncertainty.

Second, report the MRL (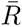) alongside statistical significance. The Rayleigh statistic 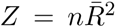 increases with observation number even when directional concentration is unchanged (Table 3B). Cells from one specimen are not independent biological replicates (Zimmerman et al.; 2021; Squair et al.; 2021). Broader conclusions require independent specimens and analyses that account for which specimen each observation comes from. Repeat the composition-before-covariance diagnostic within each specimen (Section 4.1; Supplementary Table S2).

Third, agreement between tricycle position and Seurat phase (Figure 8) can reflect their use of related cell-cycle genes. Such agreement does not independently validate the inferred positions. Biological validation should use measurements obtained separately from the expression data used to infer the angle (Section 3.3).

Fourth, check convergence before interpreting or comparing models (Mulder and Klugkist; 2017). In AD, we retained the converged projected-normal model after Bayesian von Mises regression fitted with brms failed to converge (Section 5.1). For large tissues, fitting within regions or on representative subsamples reduces computing demands. Evaluate spatial predictions on held-out cells separated from training locations, because nearby training cells can make prediction appear more accurate (Section 4.2).

## 6 Discussion

We illustrated an integrated analytical workflow for circular data analysis in single-cell and spatial omics, organized around the four considerations introduced in this paper. These considerations concern provenance and observational unit, the inferential objective, composition before covariance, and the choice of predictive or spatial circular models. The two case studies show that these considerations, rather than any single model, shape which conclusions a dataset can support.

Before any of these considerations applies, however, circularity itself must be justified, because it is a biological assumption rather than a data type. Cell-cycle position and circadian phase are periodic and polarity is directional, but an angle derived from tissue coordinates is circular only when a meaningful center, boundary, axis, or anatomical reference frame exists. Spatial omics makes this both an opportunity and a challenge, since such frames may be ambiguous or may coexist within one section, so an apparent directional pattern can reflect the chosen reference frame just as it can reflect tissue composition.

Interpretation therefore depends as much on how the angle was defined as on which model was fitted. We recommend that studies report the provenance, observational unit, reference frame, and biological replication of every circular quantity, and assess composition before modeling spatial covariance. Three directions would extend this workflow. First, uncertainty in latent angles and reference frames should be propagated into downstream inference. Second, hierarchical and scalable spatial circular models are needed for multiple specimens and region-specific covariance, which the single-specimen subsample could not address. Third, benchmarks with known circular structure would allow circular methods to be compared under realistic tissue heterogeneity. Progress on all three will require statistical methods, scalable computation, and accessible software to develop together, so that circular quantities in spatial omics can be analyzed as carefully as they are measured.

## Code and Data availability

The processed AD and ovarian-cancer tables analyzed in this study are archived at Zenodo (https://doi.org/10.5281/zenodo.23022286). These tables were derived from the spatial transcriptomic data of Gelber et al.; (2026) and the public 10x Genomics Xenium Prime FFPE human ovarian-cancer dataset (10x Genomics; 2024). The R code for reproducing the analyses is available on GitHub (https://github.com/c16267/circular_ST).

