## Supplementary note for "Circular Data Analysis for Spatial Omics"

### S1 Overview and scope

This supplement illustrates how the circular methods discussed in the manuscript are implemented in R. Short examples connect each method to its package, input format, principal function, and basic output using the AD and ovarian-cancer datasets. The complete analysis code is available at https://github.com/c16267/circular_ST and archived at Zenodo(https://zenodo.org/records/23022287).

The examples emphasize function usage. Full tuning, repeated subsampling, and manuscript-specific plotting are provided in the repository. Run the data preparation once, then select the relevant examples. Illustrative splits, sample sizes, and MCMC settings below need not reproduce the numerical results of the manuscript runs.

| Method | Package and principal functions | Example |
|---|---|---|
| Circular summaries and plots | `circular`: `circular`, `mean.circular`, `rho.circular`, `rayleigh.test`, `rose.diag`, `density.circular` | S2, S9 |
| Harmonic regression | `stats`: `lm`, `anova` | S3 |
| Random forest | `ranger`: `ranger`, `predict` | S4 |
| Circular-response network | `nnet`: `nnet`, `predict` | S5 |
| Circular-response regression | `circular`: `lm.circular`; `brms`: `brm`; `bpnreg`: `bpnr`, `coef_circ` | S6–S8 |
| Classification tree | `rpart`: `rpart`, `predict` | S10 |
| Spatial diagnostic | Base R: `dist`, `outer`, `aggregate` | S11 |
| Von Mises mixtures | `BAMBI`: `fit_angmix`, `fix_label`, `latent_allocation`; `loo`: `waic` | S12 |
| Projected spatial process | `CircSpaceTime`: `ProjSp`, `ConvCheck`, `ProjKrigSp` | S13 |

Table S1: Methods, R interfaces, and code examples. Historical supplementary results are presented alongside the corresponding applications.

### S2 Data preparation and circular summaries

#### S2.1 Load the two processed datasets

Download the two processed tables from Zenodo (https://zenodo.org/records/23022287) and set `DATA_DIR` to their local folder. Install the packages needed for the selected examples with `install.packages()`. The `brms` package additionally requires a working Stan toolchain.

The GP example uses `CircSpaceTime` 0.9.0, available as source from the CRAN archive. Package namespaces are written explicitly so that each function's source is visible.

The AD table contains derived gene-by-context phases in radians. The ovarian table contains cell-level angles, tissue coordinates in micrometres, pathology annotations, and Seurat phase labels. The code keeps descriptive and regression filters separate. For regression, `log_amp` and `amp_response` spell out the two amplitude transformations used in the repository examples.

```
DATA_DIR <- "data"  # folder containing the downloaded tables
ad <- read.csv(file.path(DATA_DIR,
  "DESeq2_region_age_gene_phase_all_genes.csv"))
ov_file <- file.path(DATA_DIR, paste0("xenium_cell_metadata_with_",
  "pathology_and_official_10x_annotation.csv.gz"))
if (!file.exists(ov_file)) ov_file <- sub("\\.gz$", "", ov_file)
oc <- read.csv(ov_file)

ad <- subset(ad, is.finite(phase_angle))
ad$phase_angle <- ad$phase_angle %% (2 * pi)
ad$genotype <- factor(ad$genotype, levels = c("WT", "APP23"))
```

```
ad$age_months <- factor(ad$age_months, levels = c(7, 14))
ad$broad_region <- factor(ad$broad_region)
ad_desc <- subset(ad, is.finite(padj) & padj <= 0.10 &
  is.finite(amplitude_log2) & amplitude_log2 >= 0.15)
cutoff <- median(ad$amplitude_log2[is.finite(ad$amplitude_log2)])
ad_reg <- subset(ad, is.finite(amplitude_log2) &
  amplitude_log2 > cutoff & amplitude_log2 > 0)
ad_reg <- droplevels(na.omit(ad_reg[, c("phase_angle",
  "amplitude_log2", "genotype", "age_months", "broad_region")]))
ad_reg$log_amp <- log2(ad_reg$amplitude_log2)
ad_reg$amp_response <- log2(1 + ad_reg$amplitude_log2)

oc <- data.frame(theta = oc$tricyclePosition %% (2 * pi),
  x = oc$x_centroid, y = oc$y_centroid,
  pathology = trimws(oc$pathology_region), phase = trimws(oc$Phase))
oc <- subset(oc, is.finite(theta) & is.finite(x) & is.finite(y) &
  !is.na(pathology) & !tolower(pathology) %in% c("", "na", "unannotated"))
```

Code S1: Read local data and prepare the variables used below.

#### S2.2 Circular objects, summaries, and visualization

`circular()` attaches angular units and conventions to a numeric vector. `mean.circular()` estimates mean direction, `rho.circular()` gives the mean resultant length (MRL), and `rayleigh.test()` tests uniformity against unimodal alternatives (see Agostinelli and Lund (2025)). The example uses AD phases; replacing `ad_desc$phase_angle` with `oc$theta` applies the same functions to ovarian cell-cycle positions. The von Mises kernel density uses an illustrative concentration bandwidth `bw = 10`.

```
th <- circular::circular(ad_desc$phase_angle,
  units = "radians", modulo = "2pi")
c(mean_rad = as.numeric(circular::mean.circular(th)) %% (2 * pi),
  MRL = circular::rho.circular(th))
circular::rayleigh.test(th)
circular::rose.diag(th, bins = 24)
plot(circular::density.circular(th, bw = 10))
```

Code S2: Circular summaries, a rose diagram, and a circular kernel density.

#### S2.3 Interpretation of circular inference

Angles are represented on $[0, 2\pi)$; signed angular differences are obtained with $\operatorname{atan2}(\sin\Delta, \cos\Delta)$. Multiplication by $24/(2\pi)$ converts AD phase differences to hours. A zero resultant has no defined mean direction, and opposing modes can produce a small MRL. Ordinary Rayleigh $p$-values assume independent observations. Pooled cells and repeated gene-by-context estimates are not independent biological replicates, so these tests are descriptive here.

| Issue | Implication | Practical response |
|---|---|---|
| Multimodality | Opposing modes can yield a small resultant and weak Rayleigh power. | Inspect the full distribution and consider broader uniformity tests. |
| Rounded or grouped angles | Ties can affect the calibration of some procedures. | Report angular resolution and use tie-aware procedures when needed . |
| Very large $n$ | Small departures can become highly significant. | Report effect size with $p$-values and use biologically interpretable units where possible. |
| Group comparison | Mean-direction and distributional differences are distinct targets. | Match the test to the target and assumptions . |
| Hierarchical sampling | Cells or spots within one specimen are not independent biological replicates. | Respect the biological sampling unit in aggregation, modeling, or permutation. |
| Model fit | A unimodal model may miss multimodal structure. | Combine goodness-of-fit assessment with graphical diagnostics and mixture models when appropriate. |

Table S2: Practical considerations for circular inference in omics data.

### S3 Alzheimer's disease analysis

#### S3.1 Provenance of the derived phase

Spot counts were aggregated into sample–region pseudobulk, and a harmonic DESeq2 model was fitted within region–genotype–age contexts (see Love et al. (2014)). With $t = 2\pi\,\mathrm{ZT}/24$, the sine and cosine coefficients $a$ and $b$ define $A = (a^2+b^2)^{1/2}$ and $\theta = \mathrm{atan2}(a, b) \bmod 2\pi$. A joint harmonic likelihood-ratio test supplies the adjusted $p$-value. These derived quantities are already present in the downloaded table, so upstream count processing is not repeated in the examples.

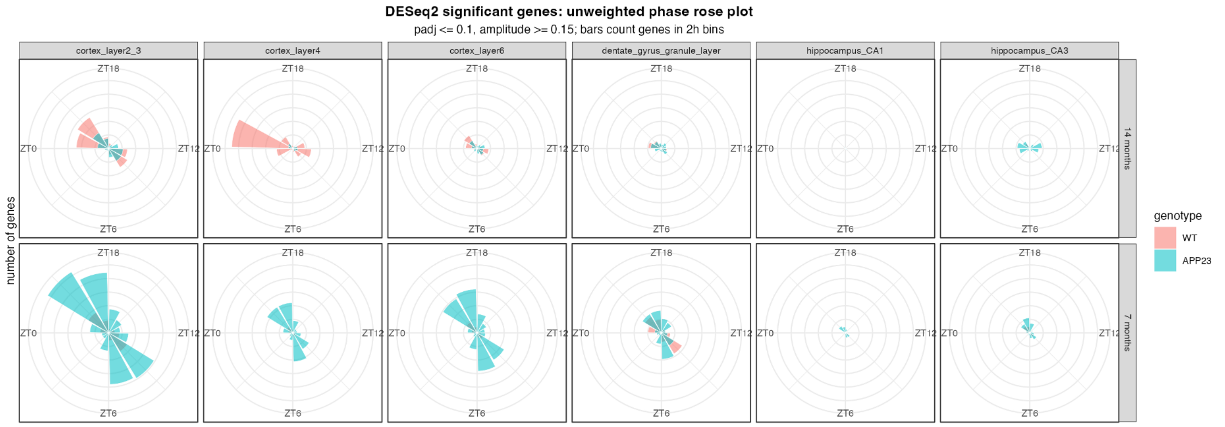

Figure S1: Fine-resolution AD peak-phase distributions for selected cortical and hippocampal annotations by genotype and age. Bars represent gene counts within two-hour phase bins.

#### S3.2 Circular description and mixture composition

Code S2 summarizes the AD descriptive set. A small pooled MRL can coexist with strong multimodality, motivating the mixture example in Code S12. Table S3 retains the manuscript composition results. Component labels are arbitrary and should be matched across fits by angular location.

| Group | $n$ | Comp. 1 | Comp. 2 | Comp. 3 |
|---|---|---|---|---|
| **Brain region** | | | | |
| Cortex | 2,232 | 13% | 50% | 37% |
| Basal ganglia | 1,071 | 18% | 47% | 35% |
| Hippocampus | 760 | 16% | 49% | 35% |
| Piriform amygdala | 265 | 25% | 48% | 27% |
| Basal forebrain and basal ganglia | 199 | 19% | 45% | 36% |
| Fiber tracts | 191 | 19% | 48% | 33% |
| **Genotype** | | | | |
| APP23 | 3,391 | 18% | 45% | 37% |
| WT | 1,609 | 13% | 55% | 32% |
| **Age** | | | | |
| 7 months | 2,936 | 13% | 47% | 40% |
| 14 months | 2,064 | 20% | 51% | 29% |

Table S3: AD composition of the three von Mises mixture components by brain region, genotype, and age. Component means are ZT13.2, ZT21.1, and ZT8.1.

##### S3.3 Phase as a circular predictor

Encode phase by its sine and cosine before supplying it to a linear or machine-learning model. In `lm()`, adding both terms gives a first-harmonic association; `anova()` compares the nested models by a joint two-degree-of-freedom test.

```
ad_reg$sin_phase <- sin(ad_reg$phase_angle)
ad_reg$cos_phase <- cos(ad_reg$phase_angle)
m0 <- stats::lm(amp_response ~ genotype + age_months + broad_region,
  data = ad_reg)
m1 <- update(m0, . ~ . + sin_phase + cos_phase)
anova(m0, m1)
plot(fitted(m1), residuals(m1)); abline(h = 0, lty = 2)
```

Code S3: Harmonic regression with context adjustment.

`ranger()` accepts the same encoded predictors and a numeric response (see Wright and Ziegler (2017)). After Code S3, the following example fits a forest on 80% of rows, predicts the held-out rows, and displays permutation importance. Phase and amplitude share their upstream harmonic coefficients, so this is an internal association and prediction exercise.

```
set.seed(123)
i <- sample.int(nrow(ad_reg), floor(0.8 * nrow(ad_reg)))
rf <- ranger::ranger(amp_response ~ sin_phase + cos_phase + genotype +
  age_months + broad_region, data = ad_reg[i, ], num.trees = 500,
  importance = "permutation", seed = 123, num.threads = 1)
pred <- predict(rf, data = ad_reg[-i, ])$predictions
y <- ad_reg$amp_response[-i]
c(RMSE = sqrt(mean((pred - y)^2)), correlation = cor(pred, y))
barplot(rf$variable.importance, horiz = TRUE, las = 1)
```

Code S4: Random-forest prediction from sine and cosine features.

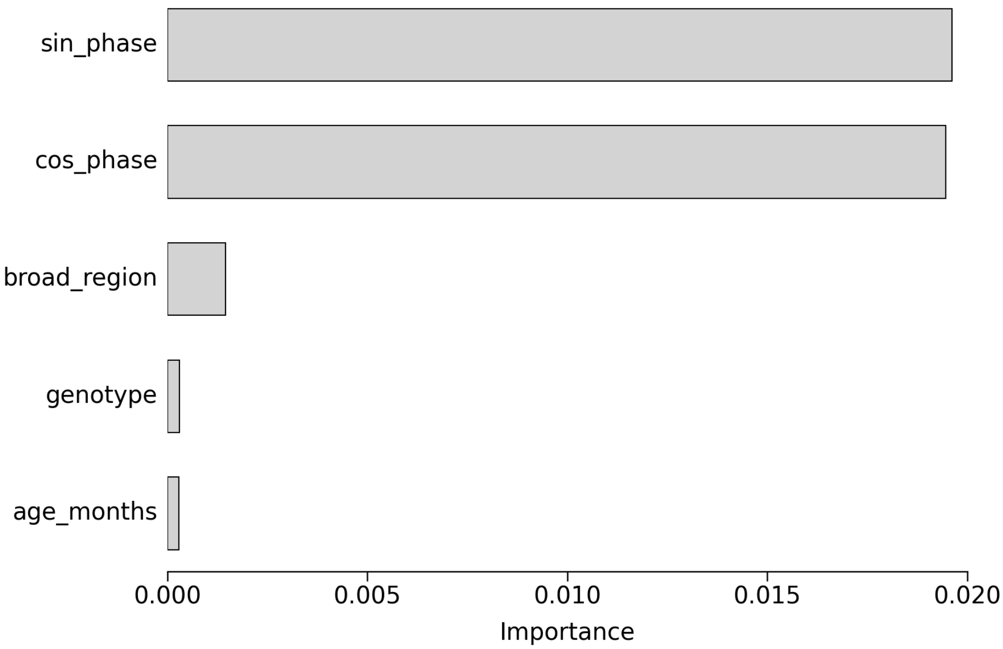

Figure S2: Permutation importance from the AD random forest predicting rhythmic amplitude. The importance values are interpreted as predictive summaries.

#### S3.4 Phase as a neural-network response

`nnet()` takes a numeric input matrix and a two-column response matrix (see Venables and Ripley (2002)). Predicting $(\sin\theta, \cos\theta)$ with linear outputs and recovering phase with `atan2()` avoids a discontinuity at $0/2\pi$. This short example fixes weight decay at 0.05 and uses a single 80:20 split; the full analysis uses training, validation, and test partitions. Scaling is learned from training rows only. The error histogram uses wrapped differences in ZT hours.

```
d <- subset(ad, is.finite(padj) & padj < 0.05 & is.finite(amplitude_log2))
d <- droplevels(na.omit(d[, c("phase_angle", "amplitude_log2",
  "genotype", "age_months", "broad_region")]))
X <- model.matrix(~ amplitude_log2 + genotype + age_months + broad_region,
  data = d)[, -1, drop = FALSE]
Y <- cbind(sin(d$phase_angle), cos(d$phase_angle))
set.seed(42)
i <- sample.int(nrow(d), floor(0.8 * nrow(d)))
r <- range(X[i, "amplitude_log2"])
X[, "amplitude_log2"] <- (X[, "amplitude_log2"] - r[1]) / max(diff(r), 1e-8)
net <- nnet::nnet(X[i, ], Y[i, ], size = 32, linout = TRUE,
  decay = 0.05, maxit = 600, MaxNWts = 10000, trace = FALSE)
u <- predict(net, X[-i, , drop = FALSE])
pred <- atan2(u[, 1], u[, 2]) %% (2 * pi)
e <- pred - d$phase_angle[-i]
e_h <- atan2(sin(e), cos(e)) * 24 / (2 * pi)
mean(abs(e_h))
hist(e_h, xlab = "Signed angular error (hours)")
```

Code S5: Two-output neural network and angular prediction error.

##### S3.5 Circular-response regression

**Frequentist von Mises regression.** `lm.circular(type = "c-l")` takes a circular response and a numeric design matrix. Its mean direction is $\mu + 2\arctan(X\beta)$, so the example removes the ordinary intercept column and supplies one initial value per slope. This benchmark is shown separately because convergence failures occurred in the AD analysis (Table S4); the function can run for a long time on unsuitable data.

```
set.seed(1)
d <- ad_reg[sample.int(nrow(ad_reg), min(500, nrow(ad_reg))), ]
X <- model.matrix(~ log_amp + genotype + age_months, d)[, -1, drop = FALSE]
vm <- circular::lm.circular(circular::circular(d$phase_angle), X,
  type = "c-l", init = rep(0, ncol(X)), tol = 1e-5)
print(vm)
barplot(vm$coefficients, names.arg = colnames(X), las = 2)
```

Code S6: A von Mises regression call on an illustrative AD subsample.

| Subsample size | 650 | 1,300 | 2,600 | 3,900 | 5,200 | 6,500 |
|---|---|---|---|---|---|---|
| Failed fits | 216 | 254 | 309 | 413 | 549 | 783 |

Table S4: Empirical stability of `lm.circular()` under repeated stratified subsampling. The second row gives the number of failed fits before ten successful fits were obtained.

**Bayesian von Mises regression.** `brm()` uses a formula and `von_mises(link = "tan_half")` (see Bürkner (2017)). Wrap the response to $[-\pi, \pi)$ before fitting. `summary()` reports posterior summaries and convergence diagnostics, and `plot()` displays traces and marginal posterior densities. The short examples use the repository's explicit log-amplitude transformation; coefficient scales must be matched when comparing fits.

```
set.seed(1)
d <- ad_reg[sample.int(nrow(ad_reg), min(500, nrow(ad_reg))), ]
d$theta <- (d$phase_angle + pi) %% (2 * pi) - pi
vm_bayes <- brms::brm(theta ~ log_amp + genotype + age_months + broad_region,
  data = d, family = brms::von_mises(link = "tan_half"),
  prior = c(brms::set_prior("normal(0,1)", class = "b"),
            brms::set_prior("gamma(2,0.5)", class = "kappa")),
  chains = 2, cores = 1, iter = 2000, seed = 1, refresh = 0)
summary(vm_bayes)
plot(vm_bayes)
```

Code S7: Bayesian von Mises regression and basic MCMC diagnostics.

**Projected-normal regression.** `bpnr()` models an angle through two latent normal regressions. The same formula is used for both components unless `pred.II` is specified (see Cremers (2024)). `fit()` returns information criteria and `coef_circ()` returns effects on the circular scale. In `bpnreg` 2.0.3, categorical contrasts are reference minus level. Only angular effects, not latent normal coefficients or predictor-axis locations, can be converted to hours.

```
set.seed(1)
d <- ad_reg[sample.int(nrow(ad_reg), min(500, nrow(ad_reg))), ]
pn0 <- bpnreg::bpnr(phase_angle ~ 1, data = d,
```

```
  its = 1000, burn = 200, n.lag = 1, seed = 1)
pn <- bpnreg::bpnr(phase_angle ~ log_amp + genotype + age_months + broad_region,
  data = d, its = 1000, burn = 200, n.lag = 1, seed = 1)
bpnreg::fit(pn0); bpnreg::fit(pn)
bpnreg::coef_circ(pn, type = "categorical")
bpnreg::coef_circ(pn, type = "continuous")
matplot(pn$beta1, type = "l", lty = 1, xlab = "Posterior draw")
```

Code S8: Projected-normal fitting, model comparison, circular effects, and traces.

Tables S4–S5 and Figure S3 retain the original diagnostic results. The reported `brms` maximum $\widehat{R} = 2.84$ precludes substantive interpretation of its information criteria. Comparisons require converged fits on the same observations within a compatible model family; trace plots alone do not establish convergence. The illustrative sample sizes and MCMC lengths above are not the original manuscript settings.

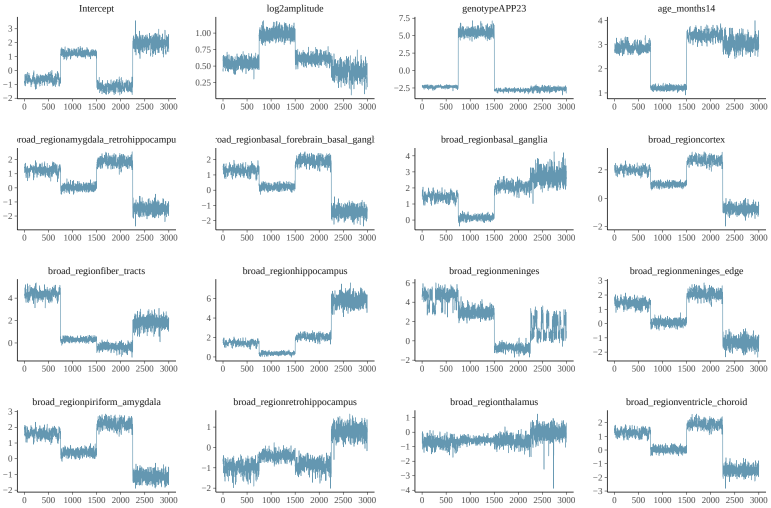

(a) `brms` von Mises model.

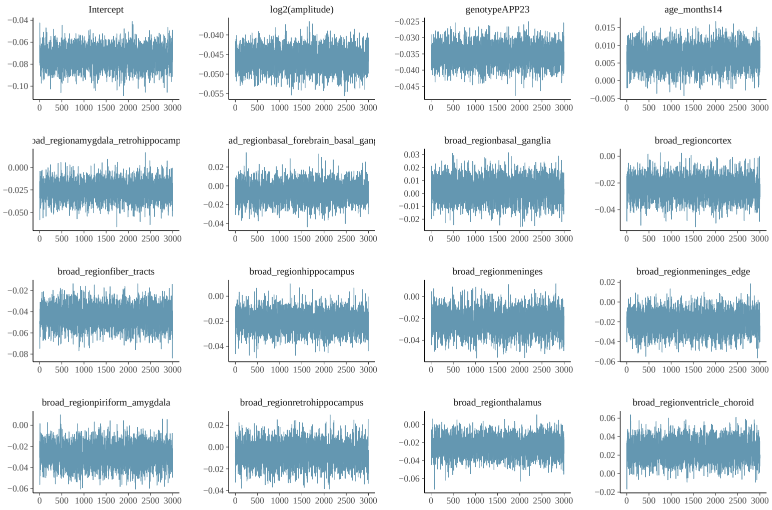

(b) `bpnreg` projected-normal model.

Figure S3: Selected MCMC diagnostics for the AD circular-response models.

| Package | Criterion | Difference |
|---|---|---|
| `brms` (von Mises) | ELPD-LOO | +88.7 |
| `brms` (von Mises) | ELPD-WAIC | +88.5 |
| `bpnreg` (projected normal) | DIC | −1674.2 |
| `bpnreg` (projected normal) | WAIC1 | −1674.1 |

Table S5: Within-model comparisons of intercept-only and predictor-inclusive Bayesian circular regressions. Differences are calculated as the predictor-inclusive model minus the intercept-only model.

### S4 Ovarian-cancer analysis

#### S4.1 Latent phase and tissue context

`tricycle` projects an expression profile onto a pretrained two-dimensional cell-cycle space and takes its direction (see Zheng et al. (2022)). The processed table already contains these angles, tissue coordinates, pathology annotations, and Seurat labels. All cells come from one specimen, so the analyses describe within-specimen heterogeneity.

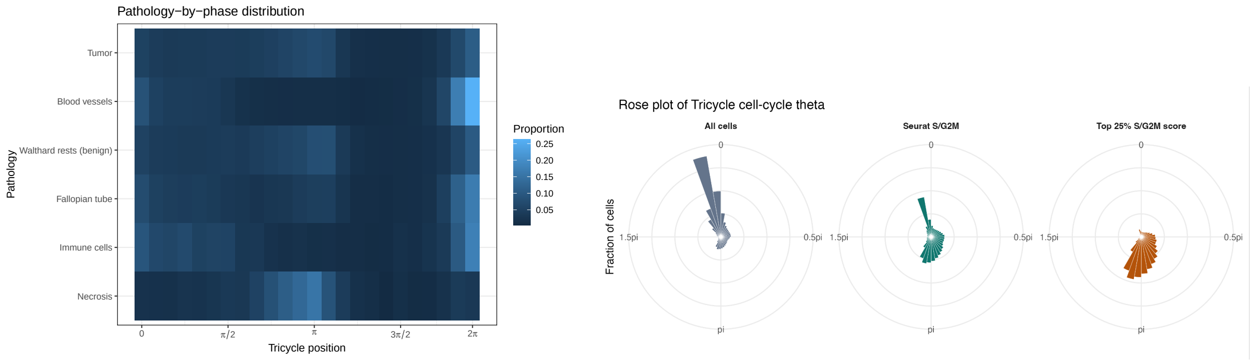

(a) Position distribution by pathology.

(b) Concordance with S and G2M information.

Figure S4: Additional context for latent cell-cycle position in the ovarian-cancer data.

#### S4.2 Pathology-specific summaries and circular clustering

Split the angle vector by pathology and apply the same `circular` functions within each group. Reporting MRL alongside the Rayleigh $p$-value is essential because $Z = n\bar{R}^2$ increases with sample size. The final call illustrates a tumor-only rose diagram; other pathology labels can be substituted directly.

```
lapply(split(oc$theta, oc$pathology), function(x) {
  z <- circular::circular(x, units = "radians", modulo = "2pi")
  c(n = length(x), mean_rad = as.numeric(circular::mean.circular(z)) %% (2*pi),
    MRL = circular::rho.circular(z), p = circular::rayleigh.test(z)$p.value)
})
circular::rose.diag(circular::circular(oc$theta[oc$pathology == "Tumor"]),
  bins = 24)
```

Code S9: Apply circular summaries within pathology groups.

The mixture interface in Code S12 applies to both applications. Table S6 retains the manuscript's three-component composition. Agreement with Seurat phase is not independent validation because both use related cell-cycle information. Biological interpretation requires matching modes to the phase reference.

| Group | $n$ | Comp. 1 | Comp. 2 | Comp. 3 |
|---|---|---|---|---|
| Seurat G1 | 2,333 | 79% | 16% | 5% |
| Seurat S | 1,469 | 28% | 48% | 24% |
| Seurat G2M | 1,176 | 15% | 8% | 77% |
| Tumor | 3,425 | 43% | 26% | 32% |
| Blood vessels | 707 | 81% | 14% | 6% |
| Walthard rests (benign) | 409 | 36% | 27% | 37% |
| Fallopian tube | 370 | 58% | 18% | 24% |
| Immune cells | 59 | 66% | 31% | 3% |
| Necrosis | 30 | 30% | 20% | 50% |

Table S6: Ovarian-cancer composition of the three von Mises mixture components. Component means are $1.95\pi$, $0.58\pi$, and $1.05\pi$.

#### S4.3 CART with a circular predictor

`rpart()` takes a factor response and the sine/cosine predictors (see Therneau and Atkinson (2025)). This example uses a split stratified by Seurat phase while retaining the observed class frequencies. Class balancing and the full validation procedure belong to the repository analysis. Mean class recall gives multiclass balanced accuracy. With `minbucket = 750`, a small dataset can produce an unsplit tree, in which case its summary is printed.

```
d <- subset(oc, phase %in% c("G1", "S", "G2M"))
d$phase <- factor(d$phase, levels = c("G1", "S", "G2M"))
d$sin_theta <- sin(d$theta); d$cos_theta <- cos(d$theta)
set.seed(1)
i <- unlist(lapply(split(seq_len(nrow(d)), d$phase), function(j)
  j[sample.int(length(j), floor(0.7 * length(j)))]))
tree <- rpart::rpart(phase ~ sin_theta + cos_theta, data = d[i, ],
  method = "class", parms = list(split = "information"),
  control = rpart::rpart.control(maxdepth = 3, minbucket = 750, cp = 0.005))
pred <- predict(tree, d[-i, ], type = "class")
cm <- table(observed = d$phase[-i], predicted = pred)
cm
mean(diag(cm) / rowSums(cm))
if (nrow(tree$frame) > 1) { plot(tree); text(tree, use.n = TRUE) } else print(tree)
```

Code S10: Classify Seurat phase with a shallow tree.

#### S4.4 Residual spatial dependence

Base R suffices for the empirical circular semivariogram: `dist()` gives tissue distances, `outer()` forms angular differences, and `aggregate()` averages $1 - \cos(\theta_i - \theta_j)$ within distance bins. There is no additional factor of one half. The dashed line is the marginal independence reference $1 - \bar{R}^2$. The same random sample supplies the pooled and tumor-only curves; the manuscript's proportional sampling is implemented in the full analysis. A sample-size cap limits the quadratic cost of pairwise distances.

```
set.seed(1)
s <- oc[sample.int(nrow(oc), min(1500, nrow(oc))), ]
```

```r
vario <- function(d) {
  D <- as.matrix(dist(d[, c("x", "y")]))
  A <- 1 - cos(outer(d$theta, d$theta, "-"))
  u <- lower.tri(D)
  b <- cut(D[u], unique(quantile(D[u], seq(0, 1, length.out = 16))),
           include.lowest = TRUE)
  v <- aggregate(cbind(distance = D[u], dissimilarity = A[u]),
                 list(bin = b), mean)
  plot(v$distance, v$dissimilarity, type = "b",
       xlab = "Distance (micrometres)", ylab = "Angular dissimilarity")
  abline(h = 1 - Mod(mean(exp(1i * d$theta)))^2, lty = 2)
  invisible(v)
}
v_all <- vario(s)
v_tumor <- vario(subset(s, pathology == "Tumor"))
```

Code S11: Distance-binned angular dissimilarity, pooled and within tumor.

Short-distance values below the reference indicate local angular similarity. Persistence within tumor cells suggests that broad pathology composition alone does not explain the pattern, but does not adjust for every biological or technical factor.

### S5 Mixture and spatial-model interfaces

#### S5.1 Von Mises mixture fitting

`BAMBI::fit_angmix()` fits a specified number of von Mises components (see Chakraborty and Wong (2021)). The example compares $K = 1, \ldots, 4$ by WAIC on one ovarian subsample. `return_llik_contri = TRUE` enables the pointwise likelihood needed by `loo::waic()`. After selecting the lowest WAIC, `fix_label()` aligns component labels, `latent_allocation()` assigns observations, and `lattice::densityplot()` dispatches to BAMBI's fitted-density plot. For AD, replace the sampling table with `ad_desc`, the angle with `phase_angle`, and the grouping variable with `broad_region`.

```r
set.seed(100)
d <- oc[sample.int(nrow(oc), min(1000, nrow(oc))), ]
fits <- lapply(1:4, function(k) BAMBI::fit_angmix(model = "vm",
  data = d$theta, ncomp = k, n.iter = 1000, n.chains = 2,
  chains_parallel = FALSE, return_llik_contri = TRUE, show.progress = FALSE))
w <- sapply(fits, function(f) loo::waic(f)$estimates["waic", "Estimate"])
cbind(K = 1:4, WAIC = w)
k <- which.min(w)
mix <- if (k > 1) BAMBI::fix_label(fits[[k]]) else fits[[k]]
BAMBI::pointest(mix)
d$cluster <- factor(BAMBI::latent_allocation(mix), levels = seq_len(k))
prop.table(table(d$pathology, d$cluster), margin = 1)
lattice::densityplot(mix)
```

Code S12: Fit, compare, allocate, and plot a von Mises mixture.

WAIC comparison and hard allocations do not replace assessment of MCMC mixing. Component integers can permute between fits; fitted component means and empirical means of allocated observations are different summaries.

#### S5.2 Projected Gaussian-process fitting and prediction

`CircSpaceTime::ProjSp()` fits a projected spatial process, and `ProjKrigSp()` predicts angles at new coordinates (see Jona Lasinio et al. (2020)). The example takes 200 cells, expresses coordinates in millimetres, and holds out the rightmost 20% by horizontal position. Priors and starting values below are illustrative and explicitly tied to this coordinate scale. `ConvCheck()` and `coda::traceplot()` inspect the chains. Predictions are generated within each chain and pooled before calculating the circular mean. Run the prediction part only after assessing convergence, extending the MCMC run as needed.

```
set.seed(1)
d <- oc[sample.int(nrow(oc), min(200, nrow(oc))), ]
xy <- as.matrix(d[, c("x", "y")]) / 1000
te <- which(xy[, 1] > quantile(xy[, 1], 0.8))
tr <- setdiff(seq_len(nrow(d)), te)
gp <- CircSpaceTime::ProjSp(x = d$theta[tr], coords = xy[tr, ],
  start = list(alpha = c(0, 0, 0.5, 0.5), tau = c(0, 0.2),
    rho = c(1, 2), sigma2 = c(1, 1.5), r = rep(1, length(tr))),
  priors = list(alpha_mu = c(0, 0), alpha_sigma = diag(10, 2),
    tau = c(-1, 1), rho = c(0.1, 10), sigma2 = c(2, 1)),
  sd_prop = list(sigma2 = 0.1, tau = 0.1, rho = 0.1,
    sdr = rep(0.05, length(tr))),
  iter = 2000, BurninThin = c(burnin = 1000, thin = 5),
  adapt_param = c(start = 1, end = 1000, exp = 0.9, sdr_update_iter = 50),
  corr_fun = "exponential", n_chains = 2, parallel = FALSE)
check <- CircSpaceTime::ConvCheck(gp, startit = 1000, thin = 5)
check$Rhat
coda::traceplot(check$mcmc)
# After checking convergence:
draws <- do.call(cbind, lapply(gp, function(chain)
  CircSpaceTime::ProjKrigSp(list(chain), coords_obs = xy[tr, ],
    coords_nobs = xy[te, ], x_obs = d$theta[tr])$Prev_out))
pred <- atan2(rowMeans(sin(draws)), rowMeans(cos(draws)))
e <- pred - d$theta[te]
e <- atan2(sin(e), cos(e))
mean(abs(e))
hist(e, xlab = "Signed angular error (radians)")
```

Code S13: Projected spatial model, convergence checks, and held-out prediction.

The original spatial illustration used a constant mean and exponential correlation, 1,199 training cells, 300 held-out cells, and four chains of 20,000 iterations with 10,000 burn-in iterations. Convergence remained incomplete ($\widehat{R} = 1.45$ for variance, 1.37 for spatial decay, and 1.18 for latent correlation). Figure S5 retains those diagnostics, so covariance parameters are not interpreted biologically. The short example demonstrates the API and a spatially separated holdout; it is not a reconstruction of that fit.

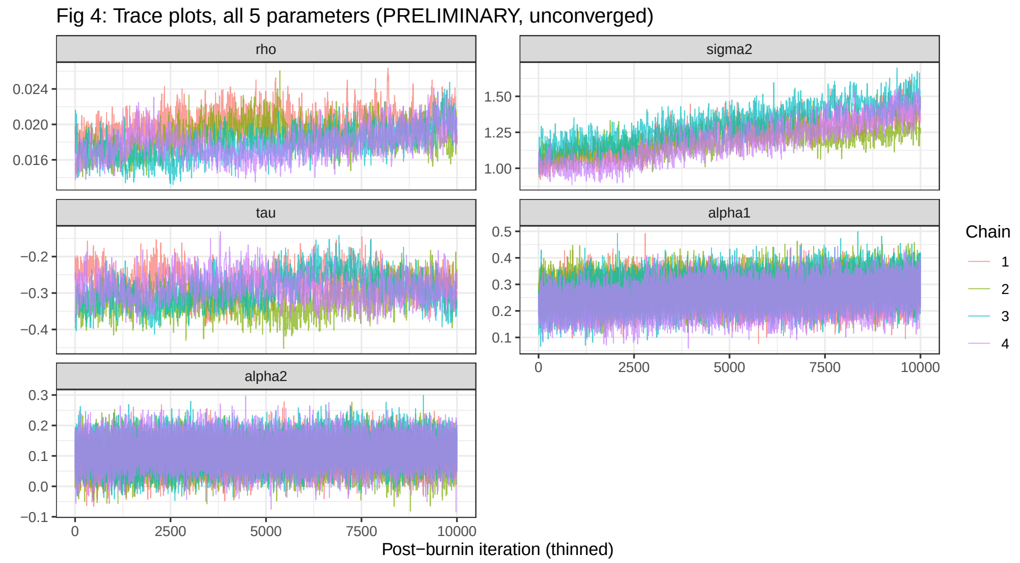

Figure S5: Projected-Gaussian-process MCMC diagnostics after burn-in. Residual drift in the variance and spatial decay parameters supports restricting interpretation to spatial prediction.
